# Role of PIF4 and FLC in controlling flowering time under daily variable temperature profile

**DOI:** 10.1101/2020.10.22.348540

**Authors:** Jutapak Jenkitkonchai, Poppy Marriott, Weibing Yang, Napaporn Sriden, Jae-Hoon Jung, Philip A Wigge, Varodom Charoensawan

## Abstract

Initiation of flowering is a crucial developmental event that requires both internal and environmental signals to determine when floral transition should occur to maximize reproductive success. Ambient temperature is one of the key environmental signals that highly influence flowering time, not only seasonally but also in the context of drastic temperature fluctuation due to global warming. Molecular mechanisms of how high or low constant temperatures affect the flowering time have been largely characterized in the model plant *Arabidopsis thaliana*; however, the effect of natural daily variable temperature outside laboratories is only partly explored. Several groups of flowering genes have been shown to play important roles in temperature responses, including two temperature-responsive transcription factors (TFs), namely PHYTOCHROME INTERACTING FACTOR 4 (PIF4) and FLOWERING LOCUS C (FLC), that act antagonistically to regulate flowering time by activating or repressing floral integrator *FLOWERING LOCUS T (FT)*. In this study, we have demonstrated that the daily variable temperature (VAR) causes early flowering in both natural accessions Col-0, C24 and their late flowering hybrid C24xCol, which carries both functional floral repressor FLC and its activator FRIGIDA (FRI), as compared to a constant temperature (CON). The loss-of-function mutation of *PIF4* exhibits later flowering in VAR, suggesting that PIF4 at least in part, contributes to acceleration of flowering in response to the daily variable temperature. We find that VAR increases *PIF4* transcription at the end of the day when temperature peaks at 32 °C. The *FT* transcription is also elevated in VAR, as compared to CON, in agreement with earlier flowering observed in VAR. In addition, VAR causes a decrease in *FLC* transcription in 4-week-old plants, and we further show that overexpression of *PIF4* can reduce *FLC* transcription, suggesting that PIF4 might also regulate *FT* indirectly through the repression of *FLC*. To further conceptualize an overall model of gene regulatory mechanisms involving *PIF4* and *FLC* in controlling flowering in response to temperature changes, we construct a co-expression – transcriptional regulatory network by combining publicly available transcriptomic data and gene regulatory interactions of our flowering genes of interest and their partners. The network model reveals the conserved and tissue-specific regulatory functions of 62 flowering-time-relating genes, namely *PIF4*, *PIF5*, *FLC*, *ELF3* and their immediate neighboring genes, which can be useful for confirming and predicting the functions and regulatory interactions between the key flowering genes.

## 1 INTRODUCTION

For the past decades, the global temperature has been fluctuating more drastically, at least in part as a result of global warming. This has major detrimental effects on all the living organisms around the world, especially plants. Since plants are sessile organisms, they are constantly exposed to environmental changes during day and night, as well as through different seasons. Thus, plants need to perceive the changes and adapt themselves to be able to respond rapidly, or face extinction. Flowering is an important developmental event in flowering plants as they need to integrate both internal and environmental factors to make an important decision on when floral transition should occur, in order to maximize reproductive success, and temperature is known to be one of the most important environmental cues that regulate flowering time (Capovilla et al., 2015; Halliday et al., 2003; Salome & McClung, 2004).

Certain plants require prolonged exposure of cold to trigger flowering, known as vernalization. The model plant *Arabidopsis thaliana* are generally classified into the winter-annual and summer-annual ecotypes, according to the time of year and how their flowering is initiated. Many winter-annual ecotypes exhibit a late-flowering phenotype that requires vernalization during the winter to trigger flowering in the following spring, while the summer-annual ecotypes can flower rapidly without the requirement of vernalization. The differences in flowering behaviors can largely be explained by the genetic variations in the genes *FLOWERING LOCUS C (FLC)* and *FRIGIDA (FRI)* (Clarke & Dean, 1994; Johanson et al., 2000; Koornneef et al., 1994; Lee et al., 1993; Michaels & Amasino, 1999; Shindo et al., 2005). Most of the late-flowering winter-annual ecotypes possess the strong (also referred to as “active” or “functional”) alleles of *FLC* and *FRI*. In contrast, most of the early-flowering summer-annual ecotypes contain either or both the weak (also known as “inactive” or “nonfunctional”) alleles of the two genes. However, strong alleles of *FLC* and *FRI* from different early-flowering plants could give rise to late flowering phenotypes when they are combined, such as in hybrid plants.

Sanda and Amasino have previously shown that the F1 plants from crossing the C24 with Col-0 ecotypes, which contain strong alleles of *FRI* and *FLC* from C24 and Col-0, respectively, flowered extremely late in relative to either parental ecotype (Sanda & Amasino, 1995). FRI, which functions as an upstream regulator of *FLC,* activates the transcription of *FLC* through forming the transcription activator complex FRI-C with the other four proteins, namely FRL1, FES1, SUF4 and FLX, to recruit chromatin modification factors and other general transcription factors (TFs) to the promoter of *FLC* (Choi et al., 2011; Michaels & Amasino, 1999). The *FLC* gene itself, which encodes a MADS-box TF, acts as a repressor of flowering by suppressing transcription of the floral integrator gene, *FLOWERING LOCUS T (FT)* (Michaels & Amasino, 1999; Searle et al., 2006). *FT* encodes a mobile signal called “florigen”, which transmits the information from leaves to shoot apical meristem to promote floral transition (Jaeger & Wigge, 2007). However, *FLC* transcription can be repressed by vernalization, which promotes the epigenetic silencing of *FLC* through the recruitment of the PHD-PRC2 complex to specific regions of *FLC* and depositing H2K27me3 repressive marks, leading to stable repression of *FLC* (Sheldon et al., 2000; Song et al., 2012). Thus, during the winter, vernalization of the winter-annual ecotypes can trigger floral transition in the following spring through epigenetic repression of *FLC* transcription.

In addition to vernalization, low and high ambient temperatures are among other temperature signals that highly influence flowering time. In laboratory conditions, constant temperatures of 22-23°C are commonly used for growing the model plant *Arabidopsis thaliana* (Rivero et al., 2014). However, the plants showed delayed and accelerated flowering when grown under a constant low temperature at 16°C, or high ambient temperature at 25°C or 27°C, respectively (Balasubramanian et al., 2006; Blazquez et al., 2003; Samach & Wigge, 2005). For an elevated temperature, it has been shown that the accelerated flowering was mainly linked to the increased expression of *FT* (Balasubramanian et al., 2006). One of the key regulators of *FT* is PHYTOCHROME-INTERACTING FACTOR 4 (PIF4), a basic helix-loop-helix (bHLH) TF, which also plays an important role in morphological changes in response to high ambient temperature, including petiole elongation, leaf hyponasty and hypocotyl elongation (Franklin et al., 2011; Gray et al., 1998; Koini et al., 2009; Sun et al., 2012). PIF4 also contributes to the induction of flowering under high ambient temperature under the short-day photoperiod (Kumar et al., 2012; Fernandez et al., 2016). However, this positive effect of PIF4 on reducing flowering time appears to be dependent on photoperiod, as the loss-of-function mutant of *PIF4* in *pif4-101*, grown at 28°C under constant light, flowered with nearly the same leaf number as in wild-type, WT (Koini et al., 2009). *PIF4*’s close homolog, *PIF5* (also known as *PIL6*), has also been shown to play a redundant role with *PIF4* in regulating hypocotyl elongation and flowering time (Fujimori et al., 2004; Kunihiro et al., 2011; Nusinow et al., 2011; Thines et al., 2014).

*PIF4* and *PIF5* are transcriptionally regulated by circadian clock (Nozue et al., 2007), where the PIF4/PIF5 proteins are degraded by the light-activated phytochrome B (phyB)-dependent mechanism (Lorrain et al., 2008; Niwa et al., 2009). During early night, the transcription of *PIF4* and *PIF5* is directly repressed by the evening complex (EC), an essential component of the circadian clock, comprising EARLY FLOWERING 3 (ELF3), EARLY FLOWERING 4 (ELF4) and LUX ARRHYTHMO (LUX), whose expressions peak at the end of day (Huang & Nusinow, 2016; Nusinow et al., 2011). Towards the end of night, the level of EC decreases, allowing the mRNA and protein of PIF4/PIF5 to accumulate, and this promotes hypocotyl growth at dawn. Interestingly, it has been shown that EC binds to its targets in a temperature-dependent manner, as the binding of EC to its targets, including *PIF4,* markedly reduces at high ambient temperature (27°C) (Box et al., 2015; Ezer et al., 2017; Silva et al., 2020). Transcript level of *PIF4* itself has been shown to increase as the temperature goes up from low to high ambient (Kumar et al., 2012; Mizuno et al., 2014), as well as its ability to bind to the *FT* promoter and activate the floral integrator gene (Kumar et al., 2012).

Despite a large body of knowledge of the molecular mechanisms of temperature-dependent flowering time control, the vast majority of the findings have been obtained in laboratory conditions using constant temperatures, or with two different temperatures during the day and night. This artificial growth environment might not be able to comprehensively represent natural conditions, where the temperature gradually rises and declines during the day and night (Wilczek et al., 2009). For this reason, Burghardt and coworkers (Burghardt et al., 2016) have adopted a variable temperature condition, where temperature profile was designed to mimic daily fluctuating temperature conditions in the real environment, and investigated its effect on flowering time. They demonstrated that this daily variable temperature condition could shorten flowering time, especially in the late-flowering plants with high *FLC* expression. In addition to *FLC*, other genes that appeared to contribute to the accelerated flowering under the variable temperature profile include *CONSTANS (CO)*, *GIGANTEA (GI), FT*, and *PHYB*, suggesting that the early flowering might not be modulated by floral repression through *FLC* alone. Interestingly, despite their well-documented functions in temperature-dependent flowering-mediating pathways, the potential involvement of PIFs in early flowering in this variable temperature condition has not been addressed. In addition, the *PIF* genes have been shown to be important for shade avoidance responses, such as hypocotyl elongation, leaf hyponasty (Lorrain et al., 2008; Nozue et al., 2015), the same phenotypes that were also strongly observed in the early flowering plants under the variable temperature condition (Burghardt et al., 2016).

In this study, we aimed to further investigate the flowering time of plants grown under the variable temperature condition that mimics daily variable temperature pattern (VAR), as used by Burghardt and colleagues (Burghardt et al., 2016), by focusing on *FLC* and *PIF4*, two key flowering genes known to be involved in temperature-dependent flowering regulation through the floral integrator *FT*. We first investigated if and to what extent, *PIF4* contributed to the acceleration of flowering under the VAR condition in natural *Arabidopsis thaliana* lines with strong and weak alleles of *FLC* and *FRI*, namely Col-0, C24, and their extremely late flowering hybrid, C24xCol-0. To gain a bigger picture of how these flowering time regulatory circuit changes with temperature, we gathered and utilized publicly available *Arabidopsis* transcriptomic data obtained from different growth temperatures and tissues, to explore transcriptional profiles of *PIF4, PIF5, FLC, ELF3,* which play a role in different but overlapping flowering pathways, as well as their immediate up- and downstream regulatory genes, under different temperature and light conditions. We also constructed a combined co-expression – transcriptional regulatory network by integrating characterized gene regulatory information with transcription patterns, which is used to investigate the dynamics and interplay among these key flowering genes in different temperatures and tissues.

## 2 MATERIALS AND METHODS

### 2.1 Plant materials and growth conditions

To investigate the potential role of *PIF4* in acceleration of flowering under the daily variable temperature condition (VAR, herein) in natural *Arabidopsis thaliana* lines with strong and weak alleles of *FLC* and *FRI*, *Arabidopsis* seeds were stratified in 0.1% ½ MS (Murashige and Skoog 1962) agar solution and kept in the dark at 4°C for 3 days to promote germination. After stratification, the seeds were sown in compost (Levington’s F2) and grown in growth chambers (MLR 351, SANYO Electric Co., Ltd.) under cool-white fluorescent light (intensity of approximately 140-160 μmol m^−2^s^−2^) in the short-day (SD) photoperiod (8 h light/16 h dark) at either constant temperature, CON (22°C), or variable temperature, VAR (22°C on average) (Figure 1a). The variable temperature profile was obtained from the previous study by Burghardt and coworkers (Burghardt et al., 2016). This part of the experiment was performed at Sainsbury Laboratory Cambridge University, UK.

**FIGURE 1.**
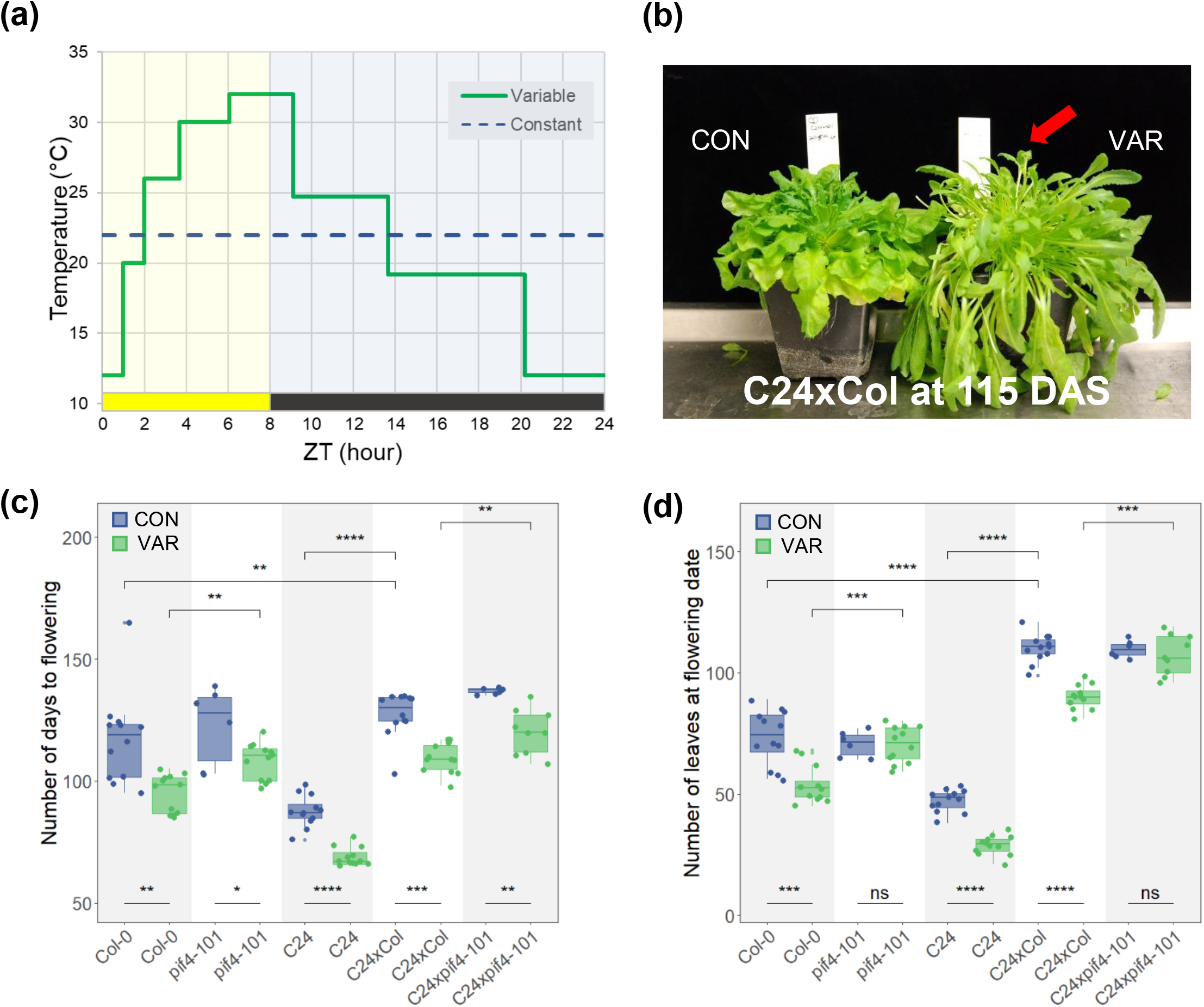
PHYTOCHROME INTERACTING FACTOR 4 (PIF4) contributes to accelerated flowering time in response to daily variable temperature. (a) The profiles of constant (CON) and variable (VAR) temperature conditions under the short-day (SD) photoperiod used in this study (adapted from Burghardt et al., 2016). CON (22°C) is shown as a dashed blue line, and VAR (also 22°C on average) is shown as a solid green line. The rectangles below the graph describe the light condition of the short-day cycle (8 h light/16 h dark), where the lights were on (yellow) and off (black); ZT, Zeitgeber time. (b) C24xCol-0 grown in CON (left) and VAR (right) conditions at 115 days after sowing (DAS). Red arrow indicates inflorescence (c) Boxplots showing the total number of days to flowering of the plants grown under the CON (blue) and VAR conditions (green). (d) Boxplots showing the total number of rosette leaves at flowering days. Solid lines inside the boxes represent medians. Each dot represents the result of each plant. There were 6 and 12 plants per line per condition. Statistical test between the two groups was computed by Wilcoxon rank sum test, p ≤ 0.0001 (****), p ≤ 0.001 (***), p ≤ 0.01 (**), p ≤ 0.05 (*) and p > 0.05 (ns).

The interplay between PIFs and their regulators, namely the evening complex (EC), and its effect on the flowering time were assessed using the mutants *pif4-101*, *elf3-2*, *pif4.pif5*, *pif4-101 elf3-2* double mutant, and *pif4.pif5 elf3-2* triple mutant, which are all in the Columbia-0 (Col-0) background. All the *Arabidopsis* seeds were gifts from the Wigge laboratory, Sainsbury Laboratory. The *pif4-101 elf3-2* double mutant and *pif4.pif5 elf3-2* triple mutant were originally generated by Nusinow and coworkers (Nusinow et al., 2011). Seeds were stratified in 0.1% agar solution in the dark at 4°C for 3 days, and sown in pots containing soil mixture (Peat moss : Perlite : Vermiculite, 3:1:1). Plants were grown on a shelf in an air-conditioned walk-in room with the light intensity of approximately 70-80 μmol m^−2^s^−2^ under cool-daylight fluorescent light (PHILIPS TL-D 36W/54-765) at a constant temperature (22±1°C) in short-day (SD) photoperiod (8 h light/16 h dark). This part of the experiment was performed at Mahidol University, Thailand.

### 2.2 Flowering time measurement

*Arabidopsis* seeds were stratified and grown under constant (CON) and variable (VAR) temperature conditions in the short-day (SD) photoperiod as described above. We measured the flowering time by the rosette leaf number and number of days to flower when the inflorescence was about 1 cm in length, as they are commonly used as flowering time indicators (Pouteau and Albertini, 2009).

### 2.3 RNA extraction and quantitative real-time PCR

Transcription levels of the flowering genes of interest, namely AT2G43010 (*PIF4)*, AT5G10140 (*FLC*) and AT1G65480 (*FT*), were assessed using RT-qPCR of the rosette leaf samples from 10-day-old or 28-day-old plants grown under the constant (CON) and variable temperature (VAR) conditions. The samples were harvested at ZT0 (dawn) and ZT8 (dusk). RNA was extracted from the leaf samples using the 96-well plate protocol as described by Box and coworkers (Box et al., 2011). Total RNA was treated with DNase I (Ambion, USA) to remove contaminating genomic DNA. DNase-treated RNA was then used for cDNA synthesis (Roche Molecular Systems, Inc.), and quantitative PCR (qPCR) was performed using SYBR Green master mix, and the Roche Lightcycler 480II instrument. The oligonucleotide primers used in this study are listed in Table S1. To measure the *PIF4* transcript levels, two primer pairs PIF4_6981/PIF4_6982, and PIF4_6581/PIF4_6582 were used to amplify different regions of the *PIF4* gene. The 6981-6982 primers amplify across the T-DNA junction, which confirms the presence or absence of T-DNA insertion at *PIF4* in the *pif4-101* line, whereas the primers 6581-6582 amplify a region upstream to the T-DNA insertion site. AT5G25760 (*PEX4*) and AT2G28390 (a *SAND family protein*) were used as reference control genes, as in previous studies (Czechowski et al., 2005; Hong et al., 2010). To investigate the interplay between PIF4/PIF5 and evening complex (EC) in regulating flowering time, transcription levels of key flowering genes, namely AT2G43010 (*PIF4*), AT3G59060 (*PIF5*) and AT1G65480 (*FT*), the samples were collected from the 28-day-old plants grown at CON (22±1°C) as described above. The aerial parts were harvested at ZT0 (dawn) and ZT8 (dusk), and were manually ground in liquid nitrogen and RNA was extracted using Trizol Reagent (Invitrogen, USA), following the manufacturer’s instruction. Removal of contaminating genomic DNA and cDNA synthesis were performed using ReverTra Ace^®^ qPCR RT Master Mix with gDNA remover (TOYOBO CO., LTD, Japan). Transcript expression was determined by qPCR using THUNDERBIRD SYBR qPCR mix (TOYOBO CO., LTD, Japan) on the Mx3000P qPCR system (Agilent technologies, USA). For this experiment, *PIF4* transcription levels were analyzed using the primers PIF4_6981/PIF4_6982. To detect *FT* transcription levels, the primers FT_7380/FT_7381 were used. AT2G28390 (a *SAND family protein*) was used as a reference control gene.

### 2.4 Data and statistical analyses

The total number of days to flowering, the total number of rosette leaves at the flowering days and transcript levels of the flowering genes of interest were displayed as box plots using the “ggplot2” package in the R statistics (R Core Team, 2019) via R studio (https://www.rstudio.com). A minimum of three biological replicates, each with three technical replicates were performed for the qPCR analysis. The relative expression was calculated using comparative Cp method (ΔΔCp). The averaged Cps of the reference genes, AT5G25760 (*PEX4*) and AT2G28390 (a *SAND family protein*) of each sample were used as controls for normalization, and the transcriptional levels from individual biological replicates were normalized again to the averaged transcriptional levels of the reference samples (normally Col-0 grown in the CON condition at ZT0, unless stated otherwise). Statistical tests between the two groups of interest were computed by Wilcoxon rank sum test using the stat_compare_means() function in the “ggpubr” package.

### 2.5 Analyzing and visualization of flowering gene transcriptomes

The list of immediate up- and downstream flowering genes of *PIF4*, *PIF5*, *ELF3* and *FLC* were extracted from the Flowering Interactive Database (FLOR-ID) (Bouche et al., 2016). Out of all 306 characterized flowering genes in FLOR-ID, our sub-network contains 62 flowering genes (*PIF4*, *PIF5*, *ELF3*, *FLC* and their 1st degree neighboring genes) (Table S2) and 128 interactions (Table S7). The transcriptional profiles of these flowering genes were extracted from 152 publicly available RNA-seq experiments of *Arabidopsis thaliana* ecotype Col-0 from 10 different studies across five different tissue types: whole seedling, rosette leaves, mature pollen grain, root and shoot apical meristem (Table S3), where the plants were treated with various temperature conditions. The RNA-seq datasets were re-analyzed and re-normalized to transcripts per million (TPM) values. To compare the transcript expression patterns between the different experimental conditions, the TPM values of each gene were Z-score transformed across either all tissues (152 samples), or across individual tissues, using the “matrixStats” package in the R statistics (R Core Team, 2019). The Z-score matrix was visualized as a heat map using the Heatmap() function in the “ComplexHeatmap” package.

### 2.6 Constructing co-expression - gene regulatory network of flowering time mechanism

The immediate up- and downstream flowering genes (1st degree neighboring genes) to the flowering-time-relating genes of interest, namely *PIF4*, *PIF5*, *ELF3*, *FLC*, were obtained from the FLOR-ID database (Bouche et al., 2016) as described above. Normalized TPM values of transcriptomes were used to calculate the correlation coefficients of transcript levels between each gene pair of 62 flowering genes across all conditions, and either across the five tissues or for individual tissues. The Spearman’s correlation coefficients were computed using the cor() function in the “ggcorrplot” package in the R statistics (R Core Team, 2019). For individual tissues, the correlation coefficient over the cutoff threshold of +/− 0.10 was considered significant, and used in further analysis. We then integrated the correlation coefficient values of individual gene pairs and the gene regulatory information from literature (i.e. activating or repressing regulatory function), to construct a combined co-expression – transcriptional regulatory network, which was visualized using Cytoscape v3.7.1 (Shannon et al., 2003).

## 3 RESULTS AND DISCUSSION

### 3.1 Daily variable temperature accelerates flowering in *Arabidopsis thaliana* Col-0, C24 and their late flowering hybrid C24xCol

*Arabidopsis thaliana* ecotypes Col-0, C24, and their late flowering hybrid, C24xCol-0 were grown at either constant temperature of 22°C (CON) or variable temperature (VAR) with the average of 22°C, as used in the earlier study by Burghardt and coworkers (Burghardt et al., 2016). The variable condition was designed to simulate the daily temperature variation, rising and falling between 12°C and 32°C, in the short-day (SD) photoperiod (Figure 1a). Phenotypic differences including leaf shape and petiole elongation were observed in the VAR condition (Figure S1). We have shown that both wild-type Col-0 and C24 grown in the VAR condition flowered earlier than those grown in the CON condition (Figure 1c). Consistently with the flowering time measured by the number of days to flowering, plants grown under VAR produced fewer rosette leaves (Figure 1d), as the plants spent less time in vegetative phase. Our second replicate (the two biological replicates were independent and were carried out separately at different times) also confirmed these trends (Figure S2). In the CON temperature condition, the F1 hybrids of C24xCol had more rosette leaves than both parental ecotypes when flowered, which was consistent with the previous study by Sanda and Amasino (Sanda & Amasino 1995). Interestingly, we observed that the delay of flowering in the hybrids were more prominent in terms of number of leaves than the number of days to flower (Figure 1d, blue boxes). As expected, C24xCol grown under VAR exhibited early flowering as compared to CON (Figure 1b, c and d), confirming that the variable temperature condition (VAR) can accelerate transition to flowering in the natural ecotypes of *Arabidopsis thaliana* Col-0, C24, as well as their late flowering hybrid, C24xCol.

### 3.2 PHYTOCHROME INTERACTING FACTOR 4 (PIF4) contributes to early flowering in daily variable temperature condition

Elevated ambient temperature is generally known to promote transition of the vegetative stage to flowering, and this phenomenon is mediated by several molecular mechanisms (see a review by Capovilla et al., 2014). In particular, previous studies revealed that accelerated flowering in warm temperature requires the activity of the bHLH TF PIF4 (Kumar et al., 2012, Thines et al., 2014). To assess whether PIF4 also contributes to earlier flowering observed in Col-0 and the late flowering hybrid C24xCol under the daily-temperature mimicking condition (VAR), we investigated the flowering time of *pif4-101* and C24x*pif4-101*, which lack two copies and one copy of functional PIF4, respectively. Under the VAR condition, the number of days to flowering of the *pif4-101* mutant was longer as compared with that of Col-0, and the difference between VAR and CON of the mutant was less than that of the wild type (Figure 1c, d). The difference between VAR and CON was even less in terms of the leaf numbers, where *pif4-101* grown in VAR flowered with nearly the same leaf number as those grown in CON (Figure 1d). This trend was also clearly observed in the second replicate in terms of leaf numbers, but less so for the number of days (Figure S2). Similarly, C24x*pif4-101* grown under VAR produced more rosette leaves and required longer time to flower than C24xCol (Figure 1c, d), suggesting the loss of functional PIF4, even in only one copy from Col-0, out of the two from each parental ecotype, partly contributed to the reduced flowering time acceleration in the VAR condition as compared to CON. However, the *pif4-101* and C24x*pif4-101* mutants grown in VAR still flowered slightly earlier than those grown in CON (Figure 1c, d and Figure S2), indicating that the variable temperature condition was still able to accelerate flowering in the full or partial absence of PIF4 activity. Thus, apart from PIF4, other regulators, including other paralogous PIF genes, might also contribute to accelerated flowering in response to the condition mimicking daily variable temperature.

### 3.3 Daily variable temperature can alter transcription levels of key flowering genes

To investigate the molecular mechanism of how PIF4 accelerates flowering time under VAR, in conjunction with the known key flowering genes of *FLC* and *FT*, the transcription levels of *PIF4*, *FLC* and *FT* were determined at 10- and 28-day after sowing (DAS), in the plants grown under CON and VAR, at two time-points: ZT0 (dawn) and ZT8 (dusk). In all the *Arabidopsis* lines tested, with an exception of *pif4-101*, *PIF4* transcription was higher in elevated ambient temperatures (Figure 2a, b), which is consistent with previous studies (Box et al., 2015; Kumar et al., 2012; Mizuno et al., 2014; Nomoto et al., 2013). At 10 DAS, *PIF4* transcription in CON (constant 22°C) was higher than in VAR (12°C at dawn, ZT0); whereas it was lower in CON (22°C) than in VAR (32°C at dusk, ZT8) (Figure 2a). The magnitudes of these differences appeared to be smaller as the plants became older (28 DAS); however, the same pattern of *PIF4* transcription remained (Figure 2b). The elevated transcription levels of *PIF4* in VAR as compared to CON at dusk, especially in the young seedlings (10 DAS), could potentially account for the early flowering seen in VAR. The might also be linked to the reduced DNA-binding activity of the evening complex (EC), hindering its repressing function of the *PIF4* expression during warm night (Box et al., 2015; Ezer et al., 2017; Nusinow et al., 2011; Silva et al., 2020). The same results were confirmed using a separate primer set (Figure S3).

**FIGURE 2.**
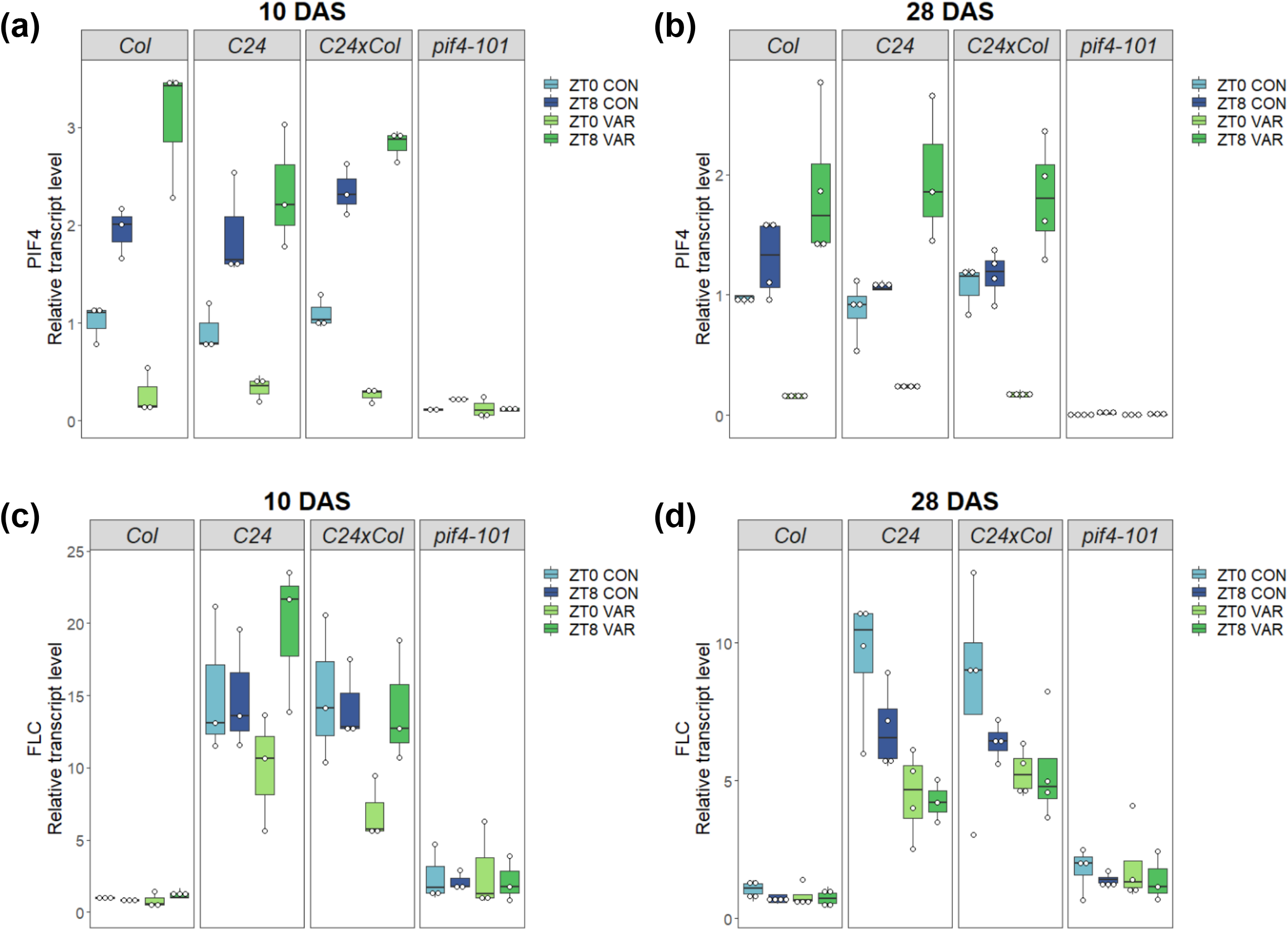
Dynamic changes of transcript levels of *PIF4* and *FLC* under the constant and variable temperature conditions. Boxplots showing the relative expressions of flowering related-genes of Col-0, C24, C24xCol and *pif4-101* of individual plants as individual white dots, normalized to the average of transcription level of Col-0 CON ZT0 (control). (a) *PIF4* at 10 DAS; (b) *PIF4* at 28 DAS (the primers used were PIF4_6981/PIF4_6982); (c) *FLC* at 10 DAS; (d) *FLC* at 28 DAS. Black lines inside the boxes represent medians.

For *FLC*, its transcription level was higher in C24 and C24xCol, where both contain a strong allele of *FRI* that can activate *FLC* expression, as compared to in Col-0 (Figure 2c, d). Interestingly, although *FLC* transcriptions appeared to be comparable at ZT0 and ZT8 in the 10 DAS plants grown at constant 22°C (Figure 2c, blue box plots), the same plants grown under VAR showed lower *FLC* transcription in the morning (ZT0) where the temperature was relatively low (12°C), and higher when the temperature was high (32°C) at dusk (ZT8) (Figure 2c, green box plots). The suppressed transcription of *FLC* in the chilly morning (12°C) of VAR might partly be due to vernalization response, as a previous study has shown that vernalization can already occur in the constant 13 °C temperature condition (Wollenberg & Amasino, 2012), although this “partial” vernalization alone could not account for the early flowering observed (Burghardt et al., 2016). At 28 DAS, the overall transcription level of *FLC* in C24 and C24xCol, as compared to that of Col-0, was reduced (from approximately 10-20 times higher than Col-0 at 10 DAS, to 5-10 times higher than Col-0 at 28 DAS) (Figure 2c, d). Importantly, *FLC* transcription of C24 and C24xCol grown under VAR was lower than CON in the 28 DAS plants at both ZT0 and ZT8, indicating that suppression of *FLC* is one potential pathway that promotes early flowering in VAR, which might be the effect of partial vernalization as described above (Figure 2d). The transcript levels of *FT*, a flowering integrator downstream to both *PIF4* and *FLC*, appeared to be increased in plants grown under the VAR condition in Col-0, C24, and their hybrid, especially at ZT8 (Figure S4), suggesting that *FT*-dependent pathway was likely to be one of the pathways that contributed to earlier flowering in response to the variable temperature. As previously reported, loss of *FT* function resulted in delayed flowering under VAR condition in L*er* but not in Col-0 background thus, this seemed to depend on the genetic background of plants (Burghardt et al., 2016). Our findings indicated that the acceleration of flowering time of plants grown in the variable temperature condition might, at least in part, be related to the suppression of *FLC* and elevated expression of the *FT* gene in VAR.

### 3.4 Overexpression of *PIF4* can reduce the effect of late-flowering phenotype under both constant and variable temperature conditions

In the variable temperature (VAR) profile, PIF4 partly contributes to accelerating floral transition, potentially through its elevated expression at the warm temperature (32°C) at the beginning of the night. This effect is especially prominent in C24xCol, the late-flowering hybrid plant carrying strong alleles of *FLC* and *FRI*. Here, we focused on the effect of elevated *PIF4* expression on the flowering time of this late-flowering hybrid, using C24 crossed to heterozygous *35S::PIF4* (in Col-0 background). A half of these F1 plants were C24x*35S::PIF4*, and the other half were C24xCol. C24x*35S::PIF4.* The plants grown in both VAR and CON flowered much earlier than C24xCol (Figure 3a), and the same trend was seen in terms of the leaf numbers (Figure 3b). However, VAR further reduced flowering time in C24x*35S:PIF4* compared to CON (Figure 3a, b). Kumar and coworkers previously found that floral acceleration by *35S::PIF4* relied on *FT* (Kumar et al., 2012). Our result showed that in terms of days to flowering, C24×35S::*PIF4* still flowered later than C24×35S::*FT* in VAR (Figure 3a), whereas the number of days to flowering were indistinguishable between C24×x*35S::FT* grown in CON and VAR (Figure 3a, b). These together indicate that apart from PIF4, other regulators also contribute to the earlier flowering time in VAR, which might also be dependent on *FT*.

We further sought to explore the mechanism of how the overexpression of *PIF4* accelerates the flowering time in the late flowering C24xCol. Transcriptional analysis by RT-qPCR showed that *PIF4* transcript level in C24x*35S::PIF4* in CON was indeed higher than that of C24xCol as expected, and also led to higher *FT* transcription (Figure 3c, e). Interestingly, we found that *FLC* transcription levels appeared to be reduced in C24x*35S::PIF4* at 10 DAS (Figure 3d) as well as at 28 DAS (Figure S5). Since *FLC* has been identified as a downstream target of PIF4 using ChIP-seq (Oh et al., 2012), it is possible that that overexpressed PIF4 binds to its cognate sites at the *FLC* promoter and suppresses its transcription. Along this line, the transcription levels of *FLC* of *pif4-101* were slightly higher than those of Col-0 in both CON and VAR (Figure 2c, d). On the other hand, the elevated expression of *PIF4* in C24xCol in VAR at ZT8 (32°C), as compared to that in CON at ZT8 (22°C) (Figure 2a), was not sufficient to suppress *FLC* within a short time frame at 10 DAS (Figure 2c), but might play a part in suppression of *FLC* over a longer period of time as seen at 28 DAS (Figure 2d).

**FIGURE 3.**
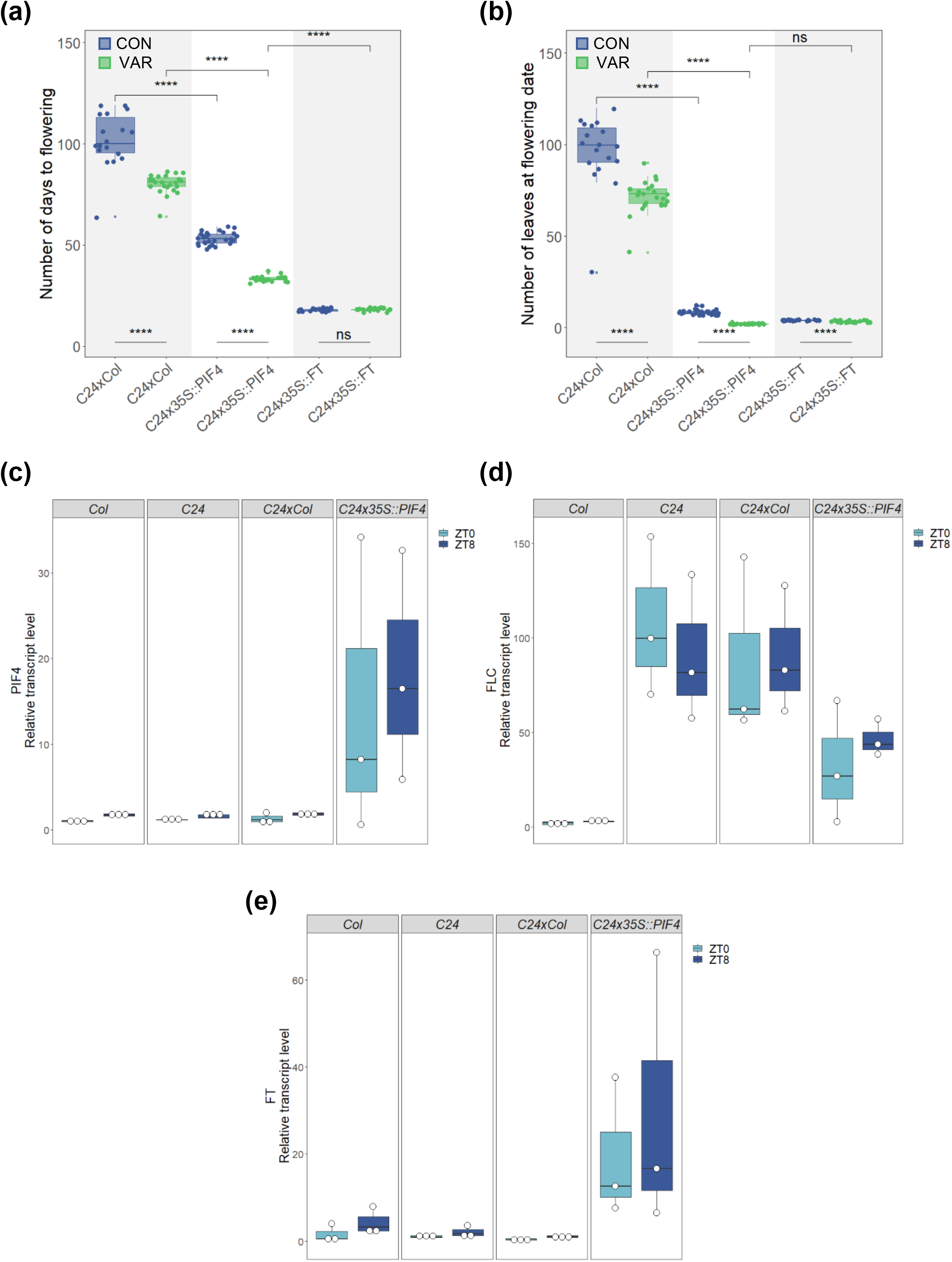
The late flowering of C24xCol can be mitigated by overexpression of *PIF4* in C24×35S::*PIF4* (a) Boxplots showing the total number of days to flowering of the plants grown under the CON (blue) and VAR (green) conditions. (b) Boxplots showing the total number of rosette leaves at the flowering days. Solid lines inside the boxes represent medians. Data points outside the boxes represent outliers. Each dot represents data from each plant. There were between 18 to 27 plants per line per condition. Statistical test between the two groups was computed by Wilcoxon rank sum test, p ≤ 0.0001 (****), p ≤ 0.001 (***), p ≤ 0.01 (**), p ≤ 0.05 (*) and p > 0.05 (ns). (c-e) Boxplots showing relative expressions of flowering related-genes namely *PIF4* (the primers used were PIF4_6581/PIF4_6582), *FLC* and *FT* (the primers used were FT_7380/FT_7381) of plants grown under constant 22°C at 10-day after sowing (10 DAS) from 3 individual plants, normalized to the average transcription level of Col-0 ZT0 (control).

### 3.5 ELF3 may bypass PIF4/PIF5 to delay flowering transition through other regulators

Having investigated the interplay between the key flowering genes *PIF4, FLC*, and *FT* and their roles in accelerating flowering time in the daily fluctuating temperature mimicking condition, here we looked to further explore the interactions of these genes and their “immediate” neighbours in the gene regulatory network. We focused on *PIF5*, a close homolog to *PIF4*, and the evening complex (EC), which directly binds to the promoters of *PIF4* and *PIF5* and represses their transcription in the early evening (Nusinow et al., 2011). It has been shown in the mutant *elf3-2*, which carries non-functional *ELF3*, a gene encoding for a component of the EC complex together with *ELF4* and *LUX*, displayed elongated hypocotyl at constant 22°C. However, this phenotype was suppressed in the *elf3-2 pif4-101* double mutant and *elf3-2 pif4.pif5* triple mutant, which showed the normal hypocotyl lengths as seen in Col-0 (Nusinow et al., 2011). Here, we asked if and to what extent the interplay between PIF4/PIF5 and the evening complex (EC) also affects the flowering time.

We grew Col-0, *pif4-101*, *elf3-2*, *pif4.pif5* double mutant, *pif4-101 elf3-2* double mutant and *pif4.pif5 elf3-2* triple mutant in constant temperature condition (22±1°C) for flowering time measurement, and transcript level analyses using RT-qPCR of the 28 DAS plants at two time-points: ZT0 and ZT8. Consistently with the results described earlier (Figure 1c, d), here we showed that loss of PIF4 function in the *pif4-101* mutant led to delayed flowering as compared to Col-0, and loss of both PIF4 and PIF5 in the *pif4.pif5* mutant led to even longer time to flower (Figure 4a) in the short day photoperiod. This indicates that PIF4 and PIF5 play a partially redundant role in flowering regulation, as previously shown by Thines and coworkers in the 12/12 h light/dark condition (Thines et al., 2014).

**FIGURE 4.**
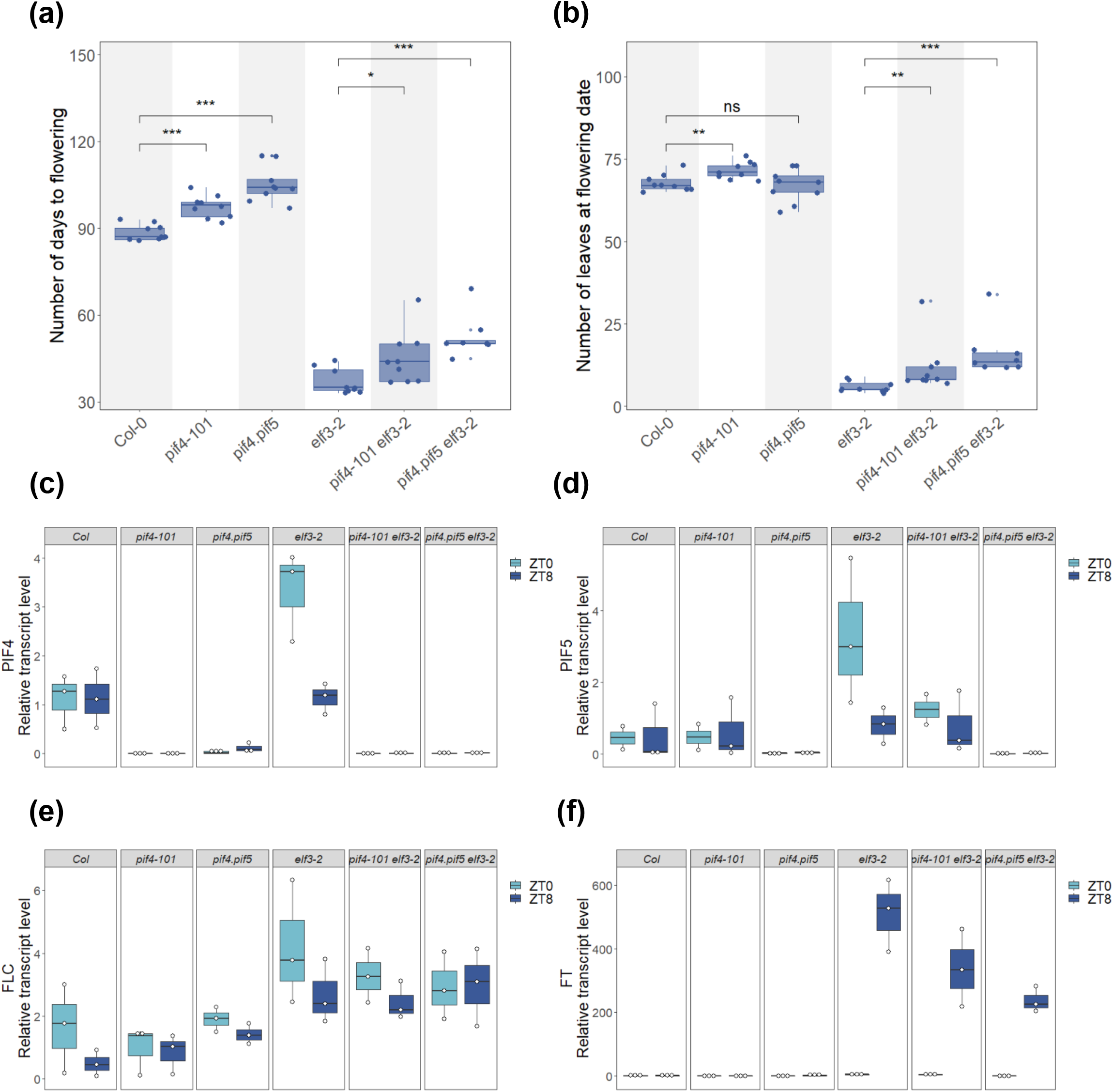
Role of evening complex (EC) in regulating flowering time through PIF4 and PIF5 and the FT pathway. (a) Boxplots showing the total number of days to flowering of the plants grown under the constant temperature condition (22±1°C). (b) Boxplots showing the total number of rosette leaves at flowering days. Solid lines inside the boxes represent medians. Data points outside the boxes represent outliers. Each dot represents data from each plant. There were 9 plants per line. Statistical test between the two groups was computed by Wilcoxon rank sum test, p ≤ 0.0001 (****), p ≤ 0.001 (***), p ≤ 0.01 (**), p ≤ 0.05 (*) and p > 0.05 (ns). (c-f) Boxplots showing the relative expressions of flowering genes namely *PIF4* (PIF4_6981/PIF4_6982 primers), *PIF5, FLC* and *FT* (FT_7380/FT_7381 primers) of plants at 28 DAS from 3 individual plants, normalized to the average of transcription level of Col-0 ZT0 (control).

The *elf3-2* mutant flowered much earlier and had fewer rosette leaves at the flowering days than Col-0 (Figure 4a, b). As the evening complex (EC) cannot be formed in the absence of ELF3 (Nusinow et al., 2011), we observed increased transcription levels of *PIF4* and *PIF5* in the *elf3-2* mutant (Figure 4c, d). The loss of either PIF4 or both PIF4 and PIF5 in the *pif4-101* and *pif4.pif5* mutants, respectively, resulted in slightly higher transcription of *FLC* as compared to Col-0 (Figure 4e) and this is similar to what was observed previously (Figure 2c). Along the same line, the loss of ELF3 in the *elf3-2* mutant showed elevated transcription of *PIF4* and *PIF5*, leading to higher transcription of *FLC* than in Col-0. However, there was no clear difference observed in the *FLC* transcription levels of the *elf3-2* mutant, the *pif4-101 elf3-2* double mutant and the *pif4.pif5 elf3-2* triple mutant, suggesting the evening complex or ELF3 alone, might also repress *FLC* transcription in an PIF4/PIF5-independent manner (Figure 4e). Interestingly, *FT* transcription levels at ZT8 were highly accumulated in the *elf3-2*, *pif4-101 elf3-2*, and *pif4.pif5 elf3-2* plants, as compared to that in Col-0 (Figure 4f). The higher *FT* accumulation could be at least in part, due to the increased expression of *PIF4* and *PIF5*, activators of *FT* transcription (Thines et al., 2014). Transcription level of *PIF5* in the *pif4-101 elf3-2* double mutant was lower than in the *elf3-2* mutant, but still higher than in Col-0 (Figure 4d). Since the previous ChIP-seq study showed that *PIF5* was a direct target of PIF4 (Oh et al., 2012), the lower transcription of *PIF5* observed in the *pif4-101 elf3-2* mutant is likely due to lack of PIF4 activity, which is partly required for transcriptional activation of *PIF5*.

Transcription level of *FT* was highest in *elf3-2*, followed by the *pif4-101 elf3-2* double mutant and *pif4.pif5 elf3-2* triple mutant (Figure 4f). However, the *FT* transcription level in the triple mutant remained higher than in Col-0. These, together with the flowering time results (Figure 4a, b and f), indicate that ELF3 (and possibly EC) regulates *FT* expression and flowering time partly through PIF4 and PIF5, but might also repress the transcription of *FT* directly or through different regulators, such as other PIF genes, resulting in the delay in transition to flowering.

### 3.6 Temperature transcriptomic profiles of flowering genes in different plant tissues

Making used of publicly available transcriptomic data, we further expanded our investigation of the transcriptional relationship among the flowering-time-relating genes of interest, namely *PIF4*, *PIF5*, *ELF3*, and *FLC*, and their immediate up- and downstream regulatory genes, based on the Flowering Interactive Database (FLOR-ID) (Bouche et al., 2016) (Table S2, see Methods), under different temperature conditions (low temperature, normal temperature, high temperature, temperature shift and heat shock), different light conditions (short-day photoperiod and long-day photoperiod) and different tissue types (whole seedling, rosette leaves, mature pollen grain, root and shoot apical meristem) (see Table S3 for complete information of transcriptomic datasets obtained). In total, we extracted 62 flowering-time-relating genes (*PIF4*, *PIF5*, *ELF3*, and *FLC* and their immediate neighbours) from the Flowering Interactive Database (FLOR-ID) (Bouche et al., 2016) (Table S2, see Methods), and their transcriptional levels from 152 RNA-seq experiments (Cortijo et al., 2017; Dickinson et al., 2018; Durufle et al., 2017; Ezer, Jung, et al., 2017; Ezer, Shepherd, et al., 2017; Martins et al., 2017; Pajoro et al., 2017; Rahmati Ishka et al., 2018; Tasset et al., 2018; Zhu et al., 2015) (Table S4).

The overall transcriptional patterns of the 62 flowering genes of interest under different temperature conditions were largely grouped by the tissue types (Figure 5a), and perhaps unsurprisingly, the transcriptional patterns of shoot apical meristem (SAM) stands out from the rest of the plant. More specifically, the genes in Cluster 3 (e.g. *ELF3*, *FCA*, *FRI*) were relatively up-regulated in SAM and root; whereas the genes in Cluster 4 (e.g. *FD*, *SOC1*, *SPL15*) appeared to be specifically up-regulated in SAM. This falls in line with earlier studies showing that *FD* is strongly expressed in SAM, and the FD protein interacts with the floral integrator *FT*, to activate the expression of another floral integrator *SOC1* in SAM, to promote floral transition (Abe et al., 2005; Searle et al., 2006; Wigge et al., 2005). SPL15 also functions together with SOC1 to activate their target genes and promote flowering transition (Hyun et al., 2016).

**FIGURE 5.**
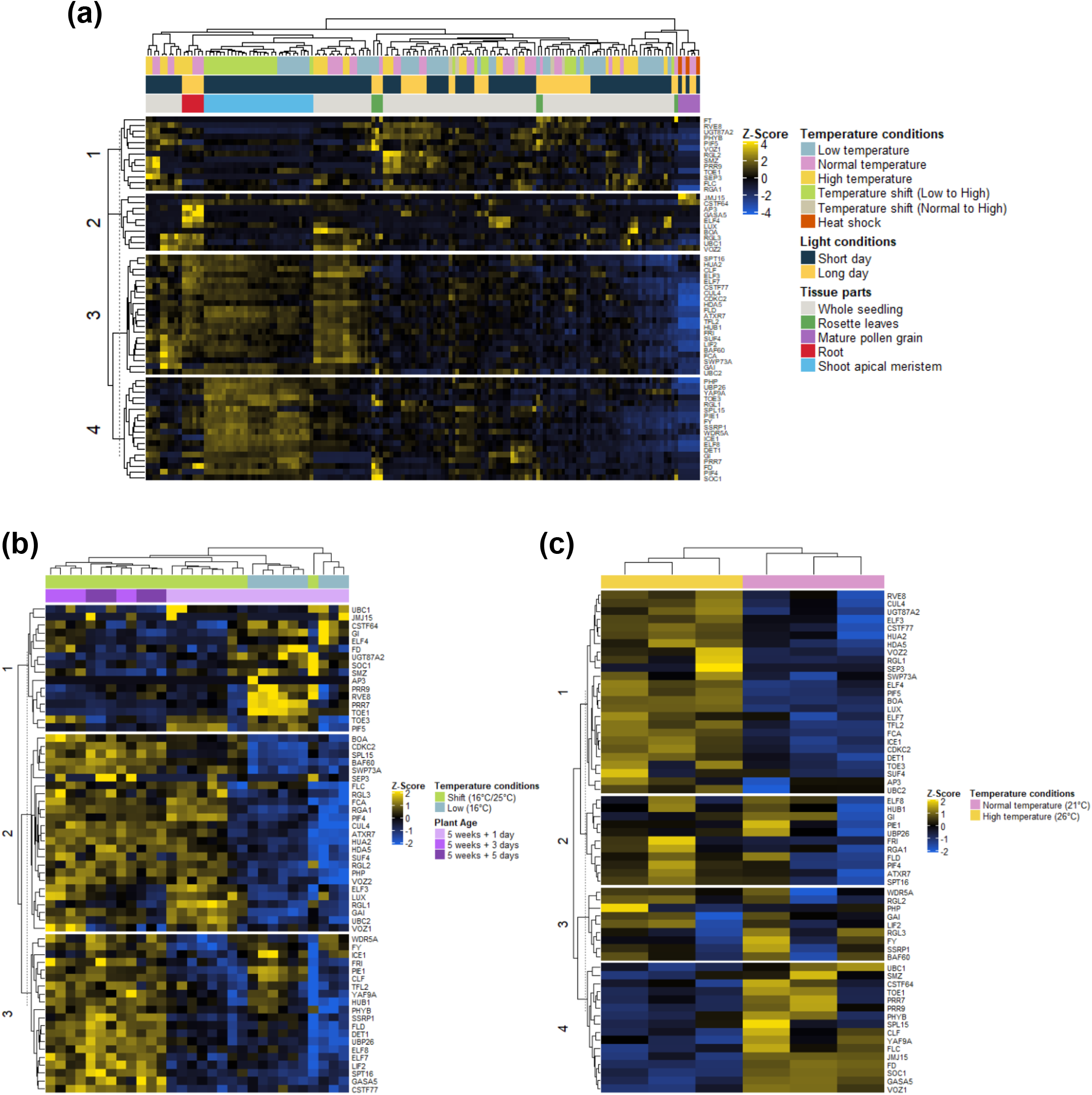
Conserved and tissue-specific transcriptional patterns of flowering genes of interest, in response to temperature changes. Heatmap showing the transcriptional patterns of 62 flowering-time-relating genes, namely *PIF4*, *PIF5*, *ELF3*, *FLC* and their immediate up- and downstream genes from (a) all five tissues, (b) shoot apical meristem, SAM (from the studies by Pajoro et al., 2017) and (c) root (from the Martins et al., 2017 study) of *Arabidopsis thaliana* ecotype Col-0 grown under different temperature and light conditions. Z-scores of TPM values were calculated across the samples. Yellow indicates relative up-regulation and blue indicates down-regulation.

We next explored the effect of the temperature conditions on these flowering genes of interest in individual tissues (Figure 5b, c and Figure S6-8). For SAM, transcriptomic patterns of these genes were extracted from 30 RNA-seq experiments published by Pajoro and coworkers (Pajoro et al., 2017) and re-normalized by us (Figure 5b). In that study, the plants were grown at 16°C for 5 weeks and either kept at 16°C, or transferred to 25°C. The genes in Cluster 1 were largely up-regulated at the constant low temperature, including *TOE1*, *GI*, *PRR7* (Figure 5b). On the other hand, Clusters 2 and 3 were both largely up-regulated in the plants shifted to 25°C, except that the Cluster 2 genes (e.g. *FCA*, *PIF4*, *RGL2*) were up-regulated within 1 day after the temperature shift and remained highly transcribed after 3 and 5 days; whereas the Cluster 3 genes (e.g. *ELF7*, *SPT16*, *GASA5*) showed slower activation after 3 days of the temperature shift. Interestingly, there are some genes in Cluster 3 that were up-regulated at low temperature and then transiently down-regulated at day 1 after the shift, and returned to the original levels within 3 days, namely *ICE1*, *PIE1* and *CLF* (see Table S5 for a complete list of genes in different clusters in SAM).

The transcriptomic patterns of these flowering genes in root were derived from the study by Martins and coworkers (Martins et al., 2017), where the plants were grown at constant 21°C or 26°C for 10 days. Interestingly, several known flowering genes were up-regulated not only in SAM but also in root. This includes *FCA*, which encodes an RNA-binding protein that acts as a positive regulator of flowering. Interestingly, it has been shown that *FCA* also plays a regulatory role in root development, and loss of its function in an *fca* mutant led to shorter root lengths and reduced numbers of lateral roots (Macknight et al., 2002). We observed that the genes in Cluster 1 of the root transcriptomic heatmap, including *FCA*, were up-regulated in response to the elevated temperature (27°C) (Figure 5c). In contrast, the genes in Cluster 4 (i.e. *FLC*, *FD*, *SOC1*) appeared to be relatively down-regulated at 27°C and up-regulated at 21°C. Despite the two studies being conducted using different temperature settings, it is still intriguing to see a number of genes being activated by higher ambient temperatures in both SAM and root (e.g. *FCA*, *LUX*, *ELF7*), as well as those appeared to be specifically activated by heat in SAM (e.g. *SPL15, GASA5, CSTF64*), or in root (e.g. *ELF4*, *UGT87A2*, *RVE8*) (see Table S6 for a complete list of genes in different clusters in root).

### 3.7 A combined co-expression - gene regulatory network dissects conserved and tissue-specific regulatory functions of key flowering genes

Finally, we asked if and how the transcriptomic profiles of the flowering genes described in the previous section could be linked to their types of regulatory functions (i.e. activator and repressor), based on the characterized interactions from the Flowering Interactive Database (FLOR-ID), as well as additional evidence from literature (see Table S7 for details). We computed the Spearman’s rank correlation coefficients, ρ (“rho”), of transcriptional patterns of the each gene pair among the 62 flowering genes of interest here, and overlaid them onto the 128 regulatory interactions connecting these genes from FLOR-ID, to construct a combined co-expression – transcriptional regulatory network (Figure 6a, and a Cytoscape session file (.cys) is also provided as an Additional File). These 128 interactions comprise activation (n=35), repression (n=67), those with unknown regulatory type (n=26). In general, one may expect the agreement between correlation coefficients and the regulatory types. For instance, the transcriptional activators and their targets are expected to be co-expressed or positively correlated; whereas the repressing gene pairs should be negatively correlated. In our combined co-expression – transcriptional regulatory network; however, these were true for a number of gene pairs in the network but not exclusively, when the transcriptomes from all the tissues available were used to compute the correlation coefficients (Figure 6a). Since we have previously found that the transcriptomic patterns of these flowering genes of interest were mostly grouped together by the tissue types rather than temperature and light conditions, it is possible that the types of regulation might be also specific to tissues. To further explore this, we re-calculated the correlation coefficients of transcription levels from each tissue separately.

**FIGURE 6.**
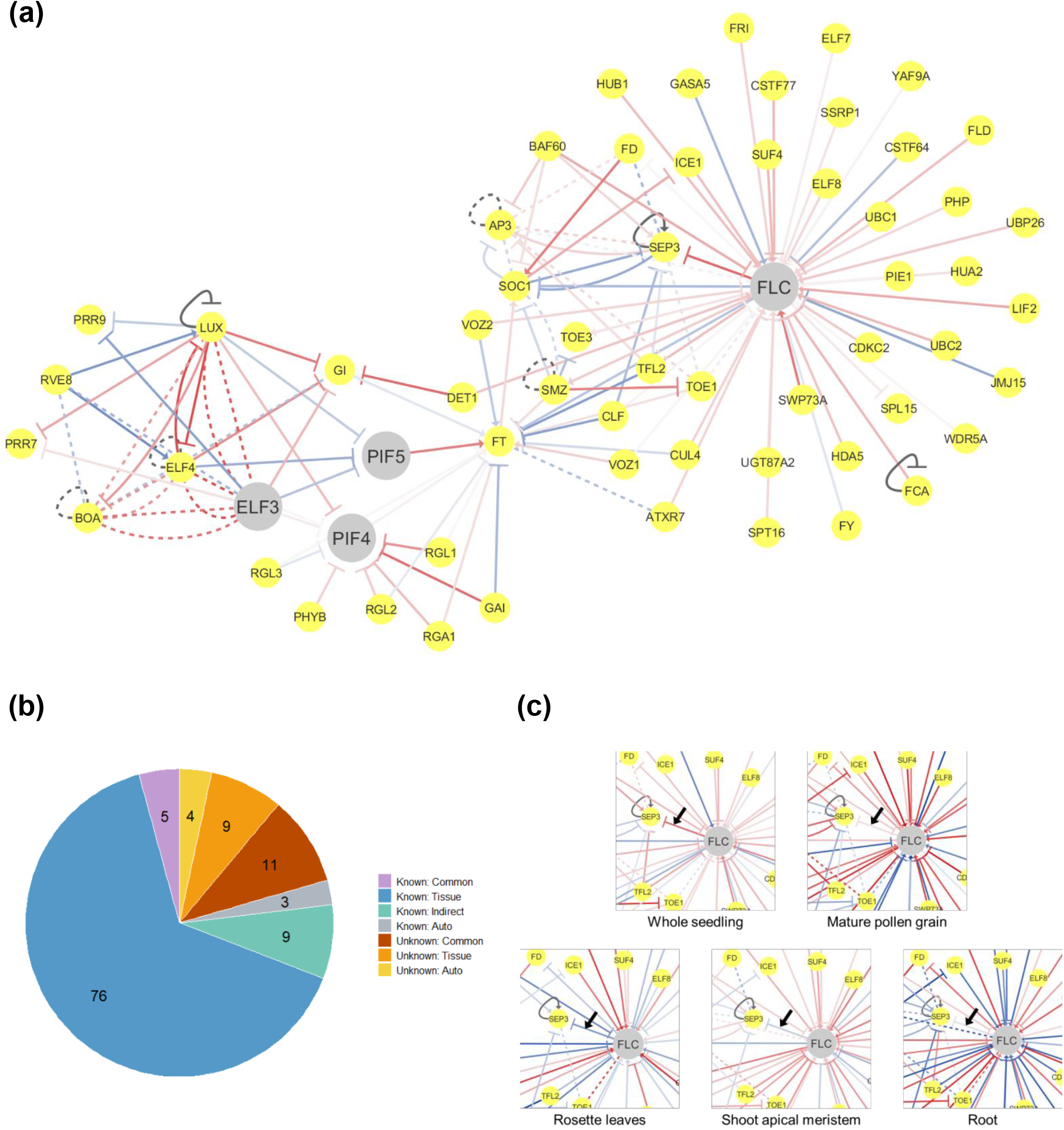
A co-expression – transcriptional regulatory network reveals the tissue-specific interactions between flowering genes. (a) Overview of the co-expression – transcriptional regulatory network of 62 flowering related-genes based on expression profiles from all available tissues. Node colors represent the flowering genes of interest, namely *PIF4*, *PIF5*, *ELF3*, *FLC* (grey) and their immediate up- and downstream flowering genes (yellow). Edge colors represent the correlation coefficients: positive (red) and negative correlation (blue), using Spearman’s correlation coefficient method. Lower correlation values are shown as paler-color edges and higher correlation values as darker-color edges. Solid/dashed black lines represent auto-regulation. (b) Pie chart showing the proportions of interaction types of gene pairs in the combined network. (c) Sub-network modules showing the expression correlations computed from individual tissues namely, whole seedling, mature pollen grain, rosette leaves, shoot apical meristem (SAM), and root. Black arrows indicate the gene pair where their co-expression coefficients vary by tissues.

Overall, our combined co-expression – transcriptional regulatory network consists of 128 interactions that could be divided into two groups: those with known (activation or repression) and unknown type of regulation (Table S7). We sub-classified these interactions based on the agreement between the previously characterized regulatory functions and the concurrence of the correlation coefficients of at least three out of five tissues, using the cutoff of |ρ| ≥ 0.10 (Figure 6b, see Methods). Out of the 128 interactions, 16 interactions showed consistent positive or negative correlation coefficients of greater than 0.10 in at least three tissues, and thus were considered as the “conserved” interactions throughout the plant tissues. Five of these common interactions were also in agreement with the known regulatory functions. For instance, *UBIQUITIN CARRIER PROTEIN 1 (UBC1)* and *FLOWERING LOCUS C (FLC)* demonstrate consistently positive correlation coefficients in whole seedlings, mature pollen grains and roots (ρ = 0.22, 0.66 and 0.66, respectively), and indeed, Gu and coworkers have shown that UBC1 activates the transcription of *FLC* through chromatin modification by monoubiquitination of histone H2B in *FLC* chromatin (Gu et al., 2009). For the 11 common interactions where their types of regulation are not known, this correlation coefficient information can potentially be used to predict the regulatory relationship between the gene pairs. For instance, it has been reported that TARGET OF EAT1 (TOE1) inhibits the activity of CONSTANS (CO), a positive regulator of *FT* transcription, and hence prevents precocious flowering (Zhang et al., 2015). *TOE1* was also identified as a target of *SEPALLATA3 (SEP3)* by ChIP-seq assay (Kaufmann et al., 2009; Pajoro et al., 2014), but the regulation type between *SEP3* and *TOE1* was not known to the best of our knowledge. Our co-expression – transcriptional regulatory network showed that the expression of *SEP3* was consistently negatively correlated with *TOE1* in rosette leaves, mature pollen grains and roots (ρ = − 0.21, −0.20 and −0.13, respectively), thus SEP3 might potentially serve as a negative regulator of *TOE1*.

Interestingly, the majority of regulatory interactions in the network (85 out of 128) showed inconsistent positive and negative correlation coefficients among the five tissues, and thus their regulation types could be considered tissue-specific. For instance, the expression of *FLC* is positively correlated with its target gene *SEP3*, a gene playing a key role in determining floral organ identity (Pelaz et al., 2000), when the expression from all the available tissues was used to compute Spearman’s rank correlation coefficient (red arrow, ρ = 0.58) (Figure 6a). However, this positive correlation would have been the opposite of what had been shown that *SEP3* is negatively regulated by FLC, where the TF directly bound to the *SEP3* promoter, and the loss of function of FLC in an *flc* mutant resulted in increased expression of *SEP3* (Deng et al., 2011). Looking into our tissue-specific co-expression – transcriptional regulatory networks, we observed that the expression of *FLC* is in fact positively correlated with that of *SEP3* only in whole seedlings and pollen grain (ρ = 0.60 and 0.13, respectively), but the two genes did show negative correlations in leaves, shoot apical meristem (SAM) and root (ρ = −0.52, −0.24 and −0.13, respectively) (Figure 6c). This suggests that FLC might repress *SEP3* in a tissue-specific manner.

Similarly, it has been showed that FRIGIDA (FRI), which can activate expression of *FLC* (Choi et al., 2011; Michaels & Amasino, 1999), is also expressed and functions in several tissues, including leaves, shoot meristem and roots, and the ectopic expression of *FRI* in those tissues led to delayed flowering. Moreover, the highest increase of *FLC* expression was observed in the accession expressing *FRI* specifically in leaves, followed by shoot meristem and roots (Kong et al., 2019). Consistent with this, our result showed strong positive correlation (ρ = 0.60) between *FRI* and *FLC* in leaves, and more moderate correlations in mature pollen grain and SAM (ρ = 0.20 and 0.27, respectively) (Figure S9). On the other hand, we found that *FRI* was negatively correlated (ρ = −0.71) with *FLC* in root (Figure S9). A possible explanation is that, *FRI* and *FLC* might play a different role in root. Indeed, it was revealed that the delay of flowering could be induced by ectopic expression of *FRI* in the root. Interestingly, the expression of *FLC* was not affected in this plant, but instead the delayed flowering might be through other activation of *FLC*-like genes, such as *MADS AFFECTING FLOWERING 4 (MAF4)* and *MAF5* (Kong et al., 2019; Searle et al., 2006).

Other interactions in the network include those with the consistent positive or negative correlation coefficients in all the tissues, but in disagreement with previously known regulatory functions (9 out of 128). For instance, P2-like TF SCHLAFMÜTZE (SMZ) that acts as a repressor of flowering and it directly binds downstream of the *FT* coding region, resulting in a repression of *FT* (Mathieu et al., 2009). In contrast, we observed that the expression of *SMZ* was positively correlated with *FT* in whole seedlings, rosette leaves and mature pollen grain (ρ = 0.11, 0.76 and 0.40, respectively). A possible explanation is that the action of SMZ might depend on other proteins or co-repressors since it has been revealed that SMZ required another floral repressor FLOWERING LOCUS M (FLM), to repress *FT* transcription, as 35S::*SMZ* plants exhibited markedly reduced *FT* expression compared with WT, but 35S::*SMZ flm* could lead to derepression of *FT* (Mathieu et al., 2009). It is possible that the TFs regulate their target genes indirectly via other TFs, and thus can be considered as “indirect” interactions. Based on information from FLOR-ID and previous publications, 7 genes were considered as “auto-regulation”, 3 out of 7 genes were self-regulated by either repressing or activating their own transcription whereas other 4 genes have been previously reported that they can bind their own genes but their regulatory functions are not known. The rest are the interactions (11 out of 128) whose regulatory functions are not known and cannot be predicted from their correlation coefficients, due to insufficient evidence of consistent coefficients above the cut-off.

## 4 CONCLUSION

The effect of high and low constant temperatures on flowering time control have been well studied to a certain extent in the model plant *Arabidopsis thaliana*. However, the effect of fluctuating temperature as in the natural daily environment is underappreciated (Burghardt et al., 2016). There are several sets of flowering genes shown to play important regulatory roles in temperature responses, in this study we mainly focused on *PHYTOCHROME INTERACTING FACTOR 4 (PIF4)* and *FLOWERING LOCUS C (FLC*), which have antagonistic roles in regulation of flowering time by activating and repressing the floral integrator gene, *FLOWERING LOCUS T (FT)*, respectively (Figure 7).

**FIGURE 7.**
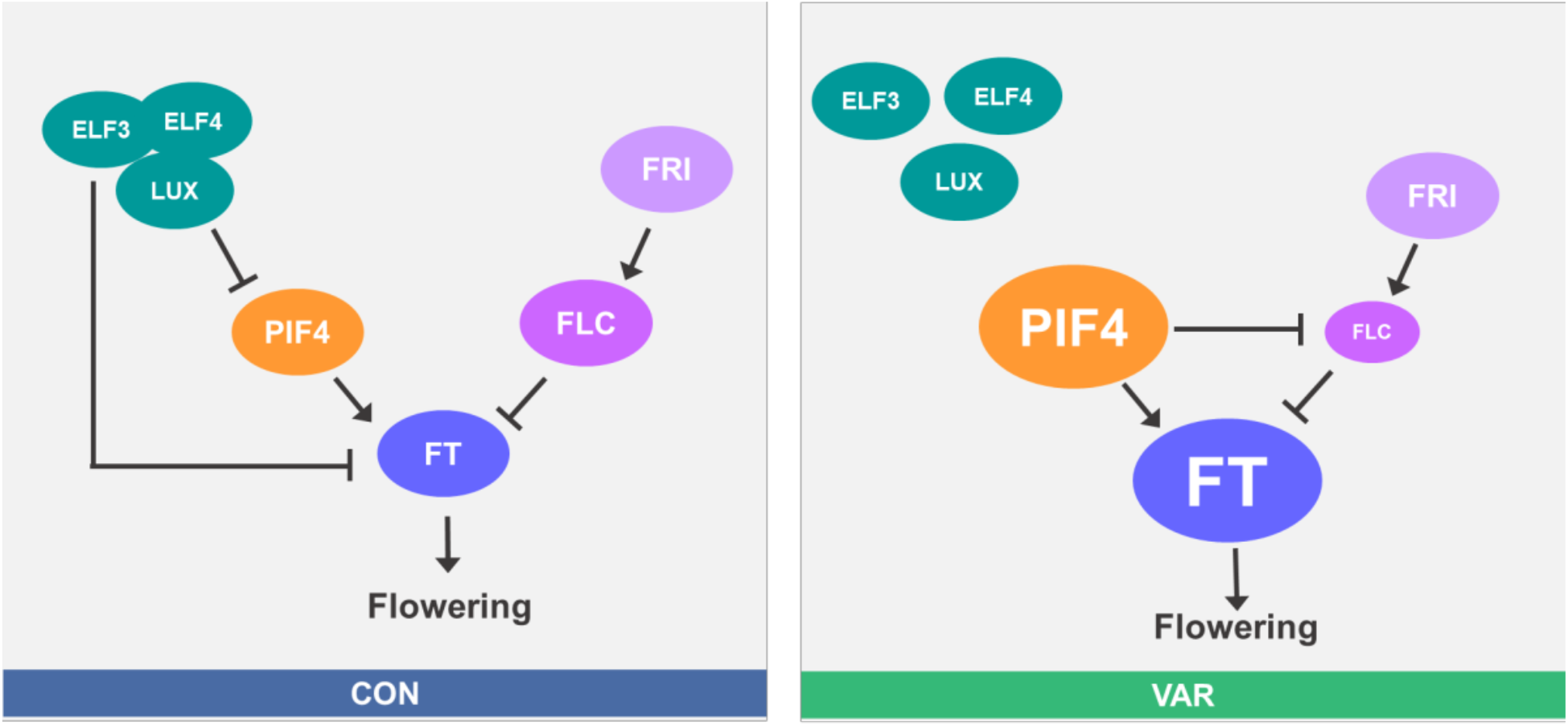
Interactions and dynamics of the flowering genes FT, PIF4, EC, FLC and FRI, and their regulatory roles in flowering time determination, in response to the daily variable temperature. The working model shows the interactions of the flowering related-genes/proteins in flowering time regulation under constant (CON) and variable (VAR) temperature conditions in the short-day (SD) photoperiod. Circles represent the genes or proteins and the sizes of circles represents the abundance (transcription levels) of genes. Solid lines with arrow and block heads indicate activation and repression, respectively.

In this study, we were able to reproduce the results showing that the daily variable temperature (VAR), which mimics a daily fluctuating temperature pattern, as used by Burghardt and colleagues (Burghardt et al., 2016), can promote the floral transition, as compared to constant temperature (CON). In addition, we have demonstrated that the phenomenon is seen in the natural *Arabidopsis* lines with weak alleles of either *FLC* or *FRI*, namely C24, Col-0; as well as their extremely late flowering hybrid, C24xCol-0, which contains the combination of strong alleles of *FLC* and *FRI* from Col-0 and C24, respectively (Figure 1c, d). *PIF4* plays an important role in acceleration of flowering time at high ambient temperature (Kumar et al., 2012); whereas the prolonged exposure to cold temperature known as vernalization can trigger the floral transition by suppression of *FLC* (Sheldon et al., 2000; Song et al., 2012) in late-flowering plants that contain both strong alleles of *FLC* and its activator *FRI* (Koornneef et al., 1994; Michaels & Amasino, 1999; Sanda & Amasino, 1995). Here, we have shown that PIF4 contributed to the early flowering in response to VAR in both natural lines and their late flowering hybrid (Figure 1c, d), and indeed *PIF4* transcription level was increased when temperature was high at dusk in VAR, as compared to CON (Figure 2a, b and Figure 7, larger circle). As shown that PIF4 contributes to the induction of flowering under a warm temperature by activating *FT* transcription (Kumar et al., 2012), here we also observed moderately elevated *FT* transcription in VAR (Figure 7, larger circle and Figure S4). Hence, we propose that, at least in part, PIF4 and FT play a role in accelerating flowering time under the daily variable temperature condition.

FLC is normally highly functional in the late flowering hybrid C24×Col (Sanda & Amasino, 1995), and here we have shown that growing the plants in VAR can reduce its transcription levels (Figure 2c, d), especially after 28 DAS. Interestingly, we noticed that the *FLC* transcription was slightly higher in *pif4-101* (Figure 2c, d), raising a possibility that PIF4 might also regulate *FT* expression and flowering time indirectly through the repression of *FLC* (Figure 7). Here, we demonstrated that the high level of *PIF4* might be sufficient to reduce the expression of *FLC* directly (Figure 2d and Figure 3d), which is in line with a previous study that found a PIF4 binding site at *FLC* promoter (Oh et al., 2012). In addition to the PIF4-dependent mechanism, the low transcription level of *FLC* was observed in VAR, also possibly due to the effect of partial vernalization (Wollenberg & Amasino, 2012), although neither partial vernalization nor high temperature alone can account for the earlier flowing time in the warm fluctuating temperatures (Burghardt et al., 2016).

Having shown the importance of PIF4 in flowering time acceleration under VAR, we looked further to the evening complex (EC), a well-characterized repressor of *PIF4*, and its potential regulatory role in this condition. We observed the increase of *PIF4* transcription level when temperature was high at dusk (Figure 2a, b), which was coincident with the reduced binding of EC to *PIF4* promoter (Box et al., 2015; Ezer et al., 2017; Silva et al., 2020), suggesting a possible function of EC in flowering time acceleration under VAR (Figure 7). However, to our surprise, our flowering time and transcription analysis results also showed that apart from *PIF4* and its homologue *PIF5*, the EC might also repress *FT* directly or through other regulators to delay flowering (Figure 4a, b, f and Figure 7). This raises a possibility that the role of EC’s in shortening the time of flowering in VAR can also be through a PIF-independent pathway. However, this is beyond the scope of this current work.

We also demonstrated a conceptual model of gene regulatory mechanisms in controlling flowering time using the combined co-expression - transcriptional regulatory network to explore the relationships between the flowering-time-relating genes of interest, namely *PIF4*, *PIF5*, *ELF3*, and *FLC*, and their immediate up- and downstream regulatory genes under different temperature conditions. This was achieved by integrating known gene regulatory functions with transcriptional expression levels from publicly available RNA-seq experiments. Based on this approach, we found that the majority of the transcriptional relationships and regulatory types between gene pairs were tissue-specific, suggesting that these flowering genes play unique regulatory roles in different tissues. This network model may help predict the regulatory functions of unknown interactions and identify the set of tissue-specific candidate genes for further exploring their roles in the flowering process. However, it should be noted that the current network model consists of 62 flowering genes, from the total of 306 flowering genes in *Arabidopsis*, characterized in the Flowering Interactive Database (FLOR-ID) (Bouche et al., 2016). In addition, more RNA-seq experiments from different plant tissues can be appended to further improve the coverage and accuracy of this network model.

Overall, we expect that this study will provide improved understanding of how plants perceive and respond to environmental conditions, such as natural fluctuation of daily temperature, in the context of flowering time, and provide a platform to breed plants that can adapt to new environments, and may help pave the way to increase crop production in the future.

## Supporting information

Additional file - Cytoscape session file

## ACKNOWLEDGEMENTS

We acknowledge the financial supports from the National Research Council of Thailand (NRCT): NRCT5-RSA63015-24 and Thailand Research Fund (TRF): MRG6080235, Faculty of Science, Mahidol University, and the Crown Property Bureau Foundation. JJ was supported by the young researcher scholarship from Faculty of Science, Mahidol University in the first year of her MSc. The authors thank Kawinnat Sue-ob for critical comments on the manuscript.

## AUTHOR CONTRIBUTIONS

Jutapak Jenkitkonchai (JJ), VC, PM, WY performed the flowering time and gene expression analyses. JJ and VC analyzed the data. JJ and NS gathered and re-analyzed the RNA-seq experiments. JJ constructed the co-expression - gene regulatory network. VC, Jaehoon Jung (JJH), PW conceived the project. JJ and VC wrote the manuscript. All authors have read and given approval to the final version of the manuscript.

## SUPPORTING INFORMATION

### Supplementary Figures

FIGURE S1 Phenotypic changes in *Arabidopsis thaliana* in response to daily variable temperature.

FIGURE S2 The second biological replicate of the flowering time results shown in Figure 1c, d. FIGURE S3 Dynamic changes of transcript levels of *PIF4* under the constant and variable temperature conditions.

FIGURE S4 Dynamic changes of transcript levels of *FT* under the constant and variable temperature conditions.

FIGURE S5 Overexpression of *PIF4* resulted in the elevated transcription levels of *FT* in older plants (28 DAS).

FIGURE S6 Transcriptional patterns of flowering genes in whole seedlings. FIGURE S7 Transcriptional patterns of flowering genes in rosette leaves. FIGURE S8 Transcriptional patterns of flowering genes in mature pollen grain.

FIGURE S9 The FRI might positively regulate the *FLC* expression in tissue-specific manner.

### Supplementary Tables

**Table S1** Oligonucleotide primers used in this study

**Table S2** A list of 62 flowering-related genes including *PIF4, PIF5, ELF3, FLC* and their up- and downstream genes

**Table S3** RNA-seq datasets and the experimental conditions used in this study

**Table S4** Normalized TPM values of 62 flowering-related genes from 152 RNA-seq datasets

**Table S5** A list of flowering-related genes in each clusters for shoot apical meristem (SAM) tissue

**Table S6** A list of flowering-related genes in each clusters for root tissue

**Table S7** The summary of the correlation coefficients in each tissues and regulatory functions of 128 interactions

### Additional File

The combined co-expression – transcriptional regulatory networks of all tissues, whole seedlings, rosette leaves, shoot apical meristem (SAM), root and mature pollen grain in Cytoscape session file (.cys) format.

Supplementary Figures

**FIGURE S1.**
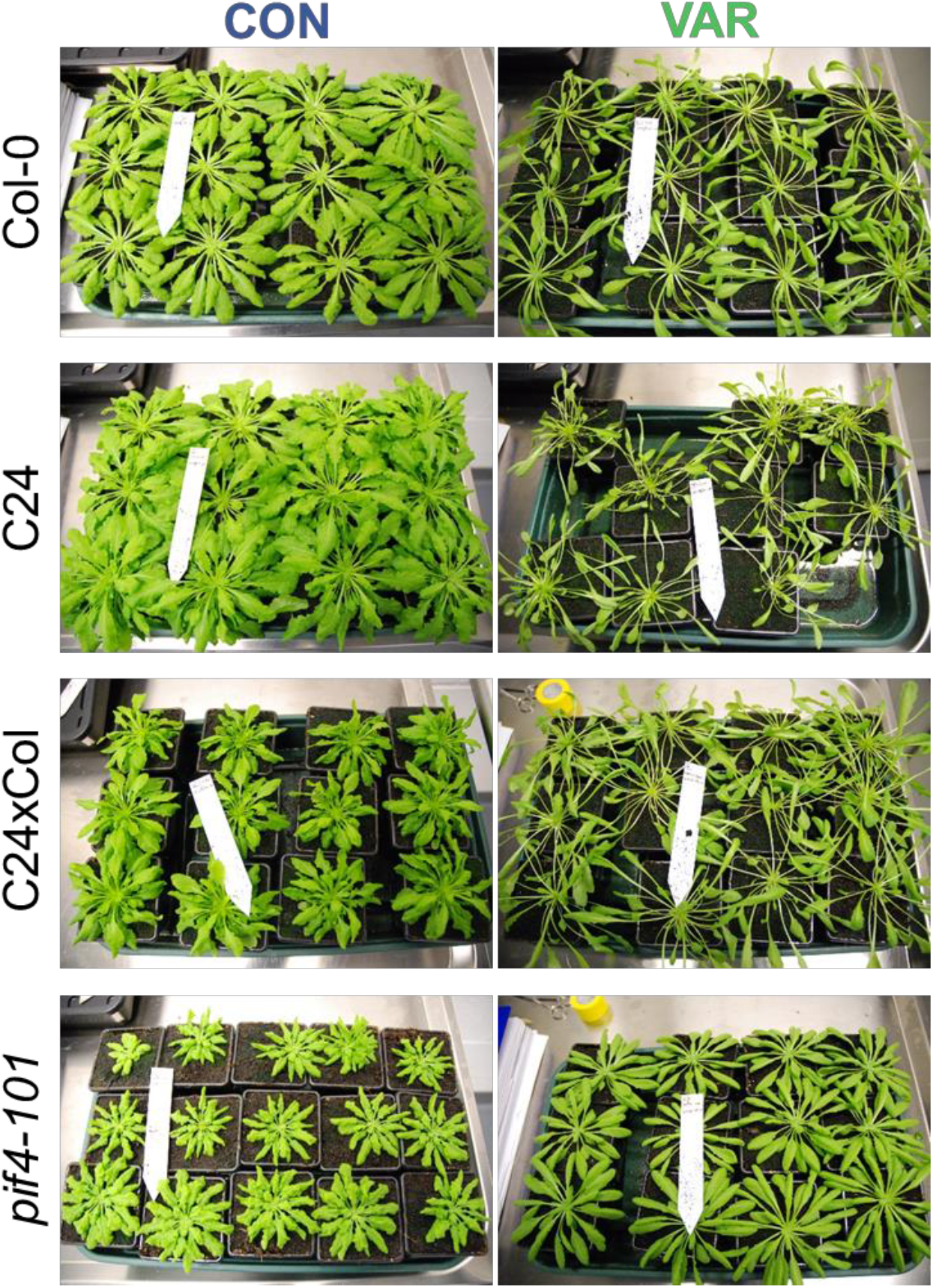
Phenotypic changes in *Arabidopsis thaliana* in response to daily variable temperature. The Col-0, C24, C24xCol and *pif4-101* plants grown at either constant temperature of 22°C (CON) or variable temperature (VAR) under the short-day (SD) condition at 70 days after sowing.

**FIGURE S2.**
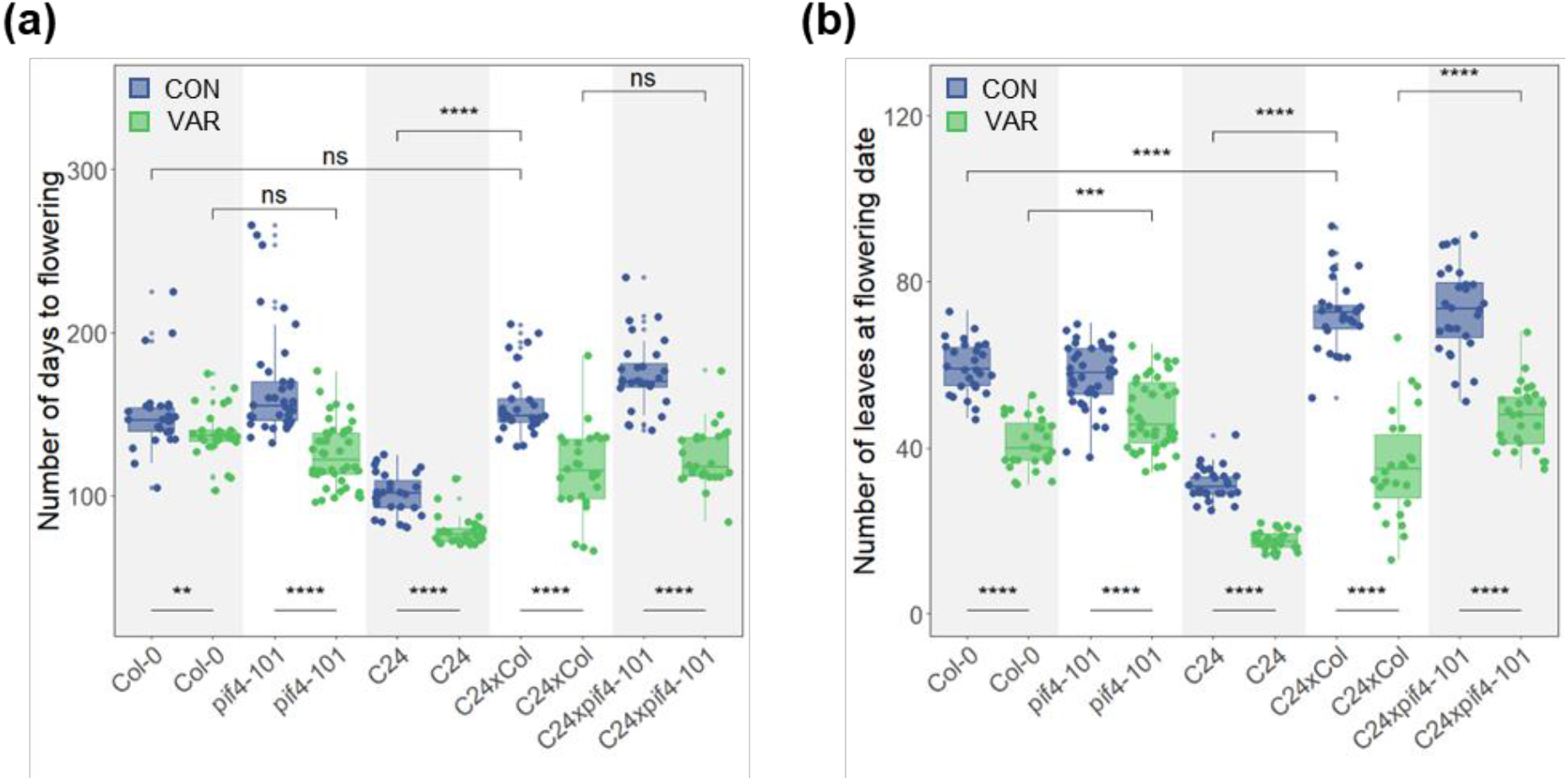
The second biological replicate of the flowering time results shown in Figure 1c, d. (a) Boxplots showing the total number of days to flowering of the plants grown under the CON (Blue) and VAR conditions (Green). (b) Boxplots showing the total number of rosette leaves at flowering days. Solid lines inside the boxes represent medians. Each dot represents data from each plant. There were 26 and 42 plants per line per condition. Statistical test between the two groups was computed by Wilcoxon rank sum test, p s; 0.0001 (****), p s; 0.001 (***), p s; 0.01 (**), p s; 0.05 (*) and p > 0.05 (ns).

**FIGURE S3.**
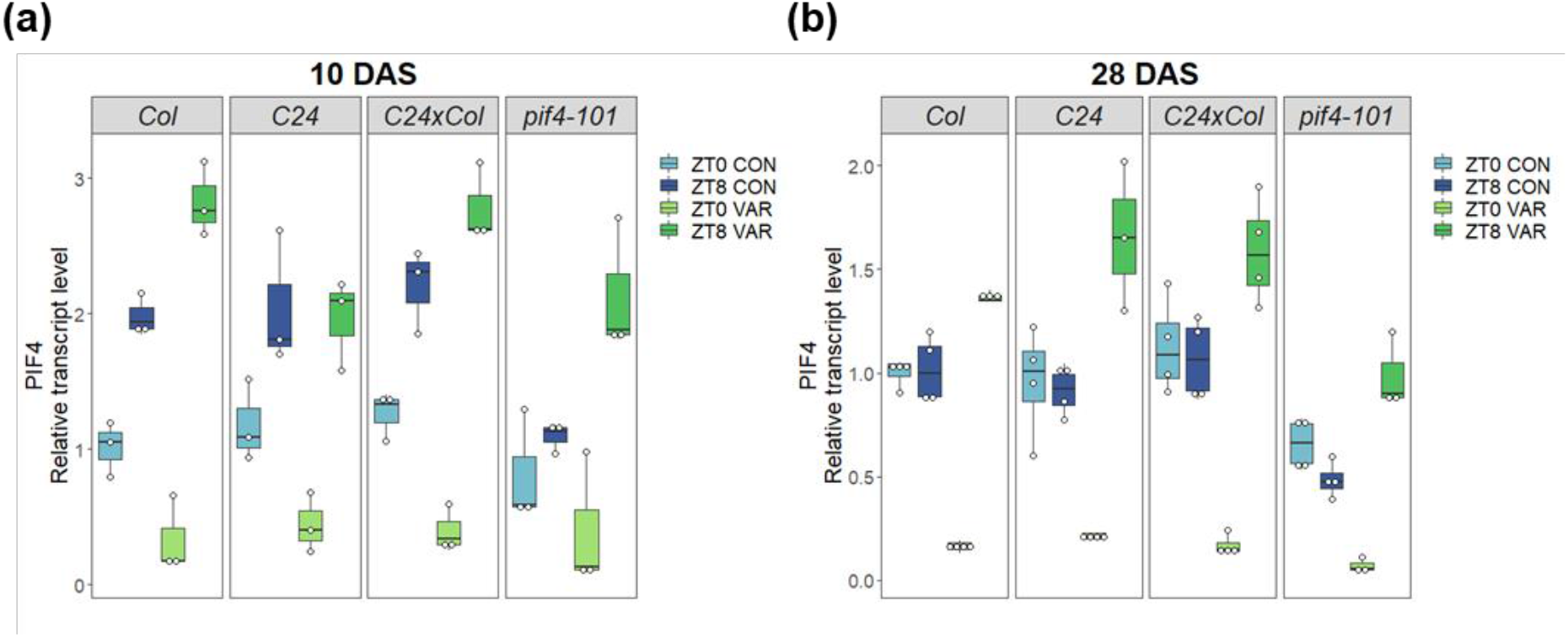
Dynamic changes of transcript levels of *PIF4* under the constant and variable temperature conditions. Boxplots showing the relative expressions of *PIF4* of Col-0, C24, C24xCol and *pif4-101* from 3-4 individual plants normalized to the average of transcription level of Col-0 CON ZT0 (control). (a) *PIF4* at 10 DAS and (b) *PIF4* at 28 DAS. The primers used were PIF4_6581/PIF4_6582. These primers were designed to amplify the sequence upstream to the position of T-DNA insertion in the *pif4-101* mutant line. White dots represent different individual plants. Black lines inside the boxes represent medians.

**FIGURE S4.**
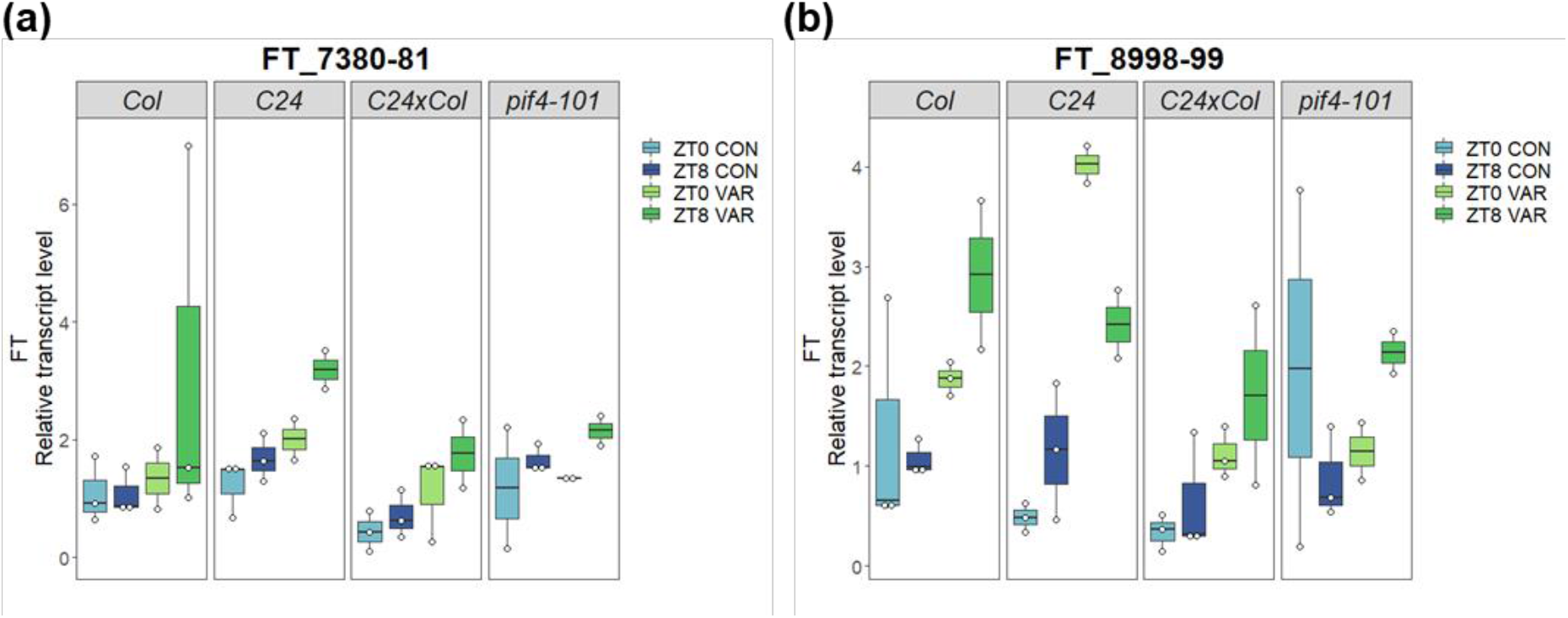
Dynamic changes of transcript levels of *FT* under the constant and variable temperature conditions. Boxplots showing the relative expressions of *FT* at 10 DAS of Col-0, C24, C24xCol and *pif4-101* from 2-4 individual plants normalized to the average of transcription level of Col-0 CON ZT0 (control). The primers used were (a) FT_7380/FT_7381 and (b) FT_8998/FT_8999. White dots represent different individual plants. Black lines inside the boxes represent medians.

**FIGURE S5.**
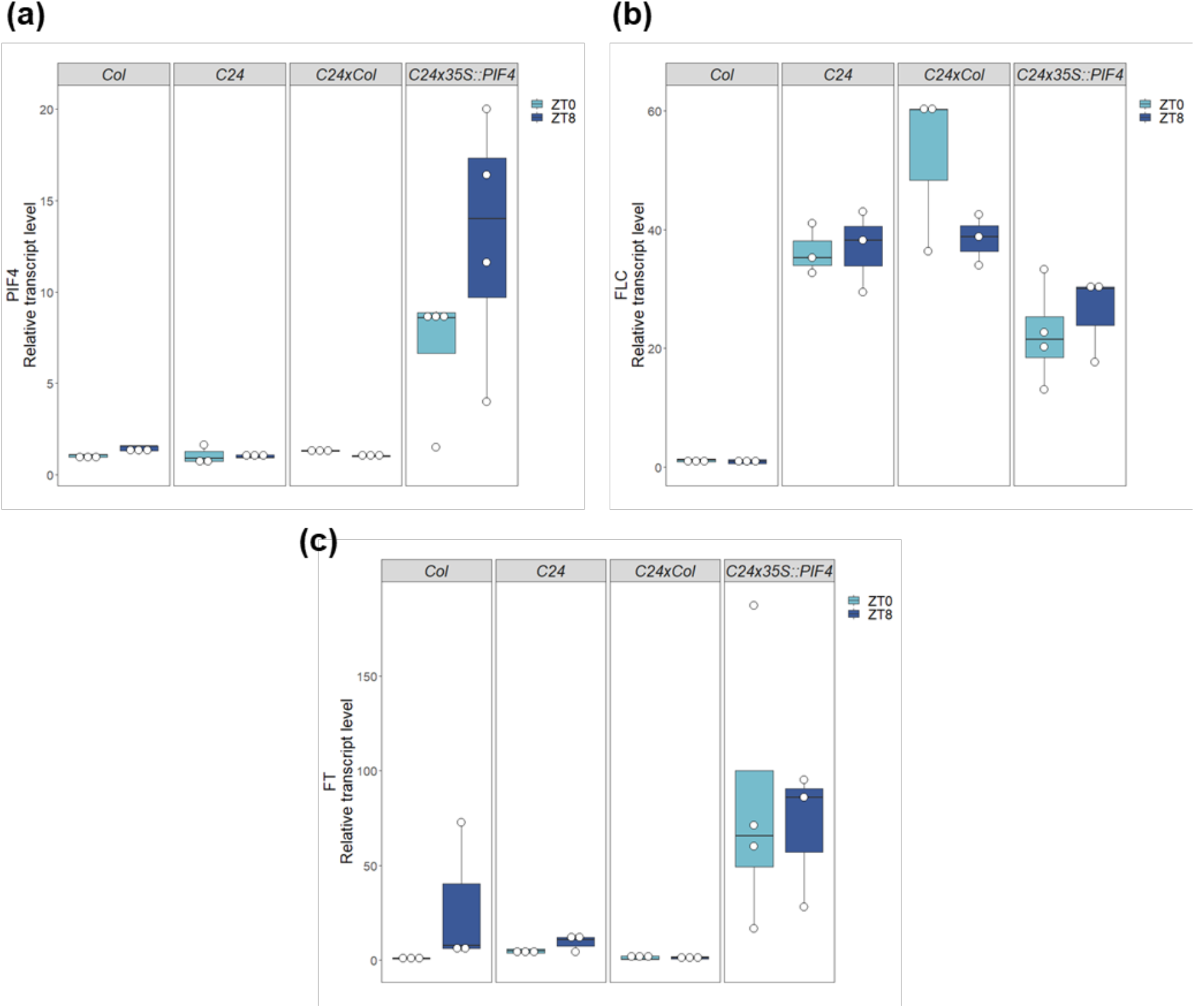
Overexpression of *PIF4* resulted in the elevated transcription levels of *FT* in older plants (28 DAS). (a-c) Boxplots showing relative expressions of flowering related-genes namely, *PIF4* (the primers used were PIF4_6581/PIF4_6582), *FLC* and *FT* (the primers used were FT_7380/FT_7381) of plants grown under constant 22°C at 28-day after sowing (28 DAS) from 3 individual plants, normalized to the average transcription level of Col-0 ZT0 (control). White dots represent different individual plants.

**FIGURE S6.**
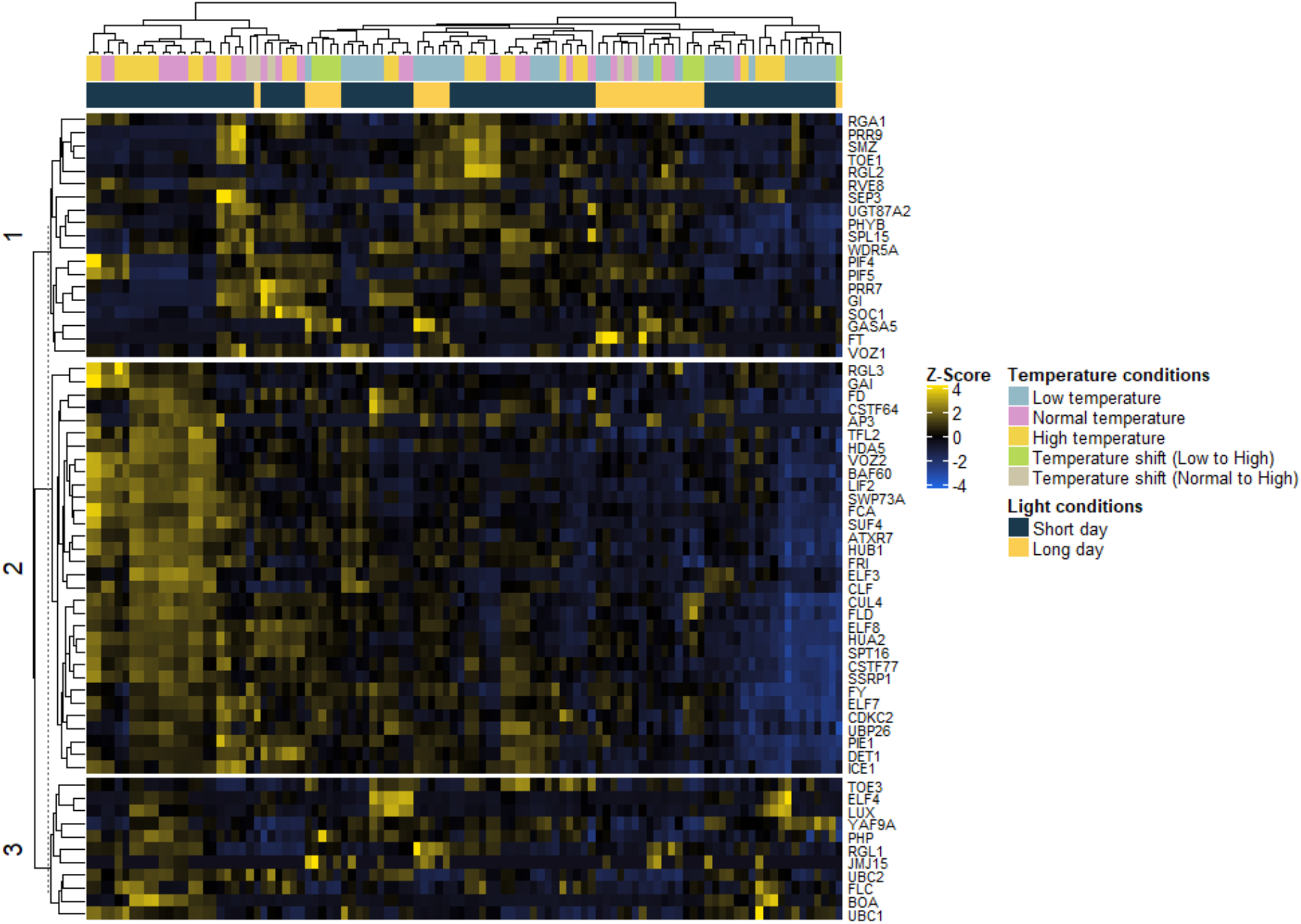
Transcriptional patterns of flowering genes in whole seedlings. Heatmap showing the transcriptional patterns of 62 flowering-time-relating genes, namely *PIF4*, *PIF5*, *ELF3*, *FLC* and their immediate up- and downstream genes from whole seedlings of *Arabidopsis thaliana* ecotype Col-0 grown under different temperature and light conditions (from the studies by Cortijo et al., 2017, Dickinson et al., 2018, Tasset et al., 2018, Zhu et al., 2015, Ezer, Shepherd, et al., 2017 and Ezer, Jung, et al., 2017). Z-scores of TPM values were calculated across the samples. Yellow indicates relative up-regulation and blue indicates down-regulation.

**FIGURE S7.**
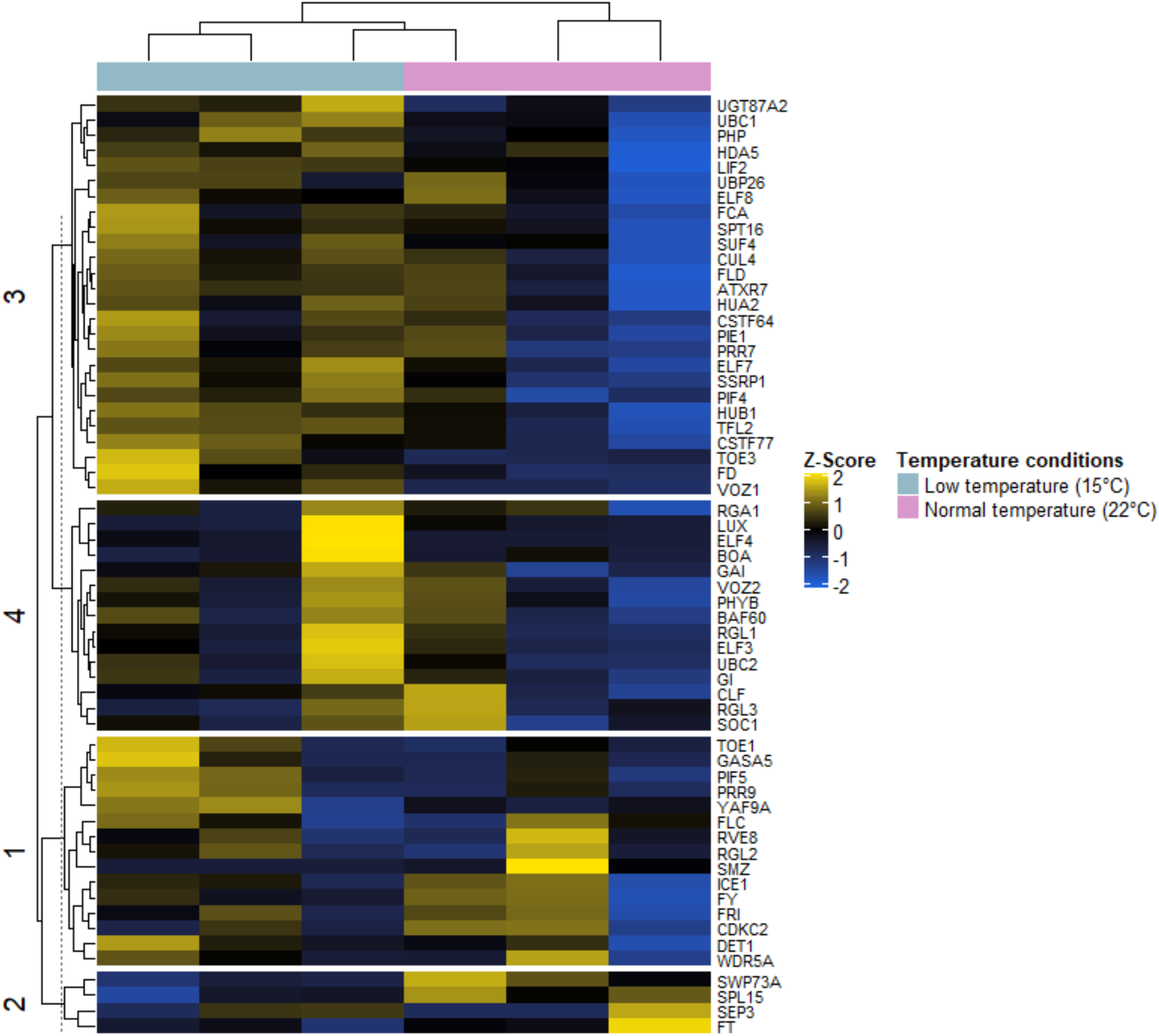
Transcriptional patterns of flowering genes in rosette leaves. Heatmap showing the transcriptional patterns of 62 flowering-time-relating genes, namely *PIF4*, *PIF5*, *ELF3*, *FLC* and their immediate up- and downstream genes from rosette leaves of *Arabidopsis thaliana* ecotype Col-0 grown under different temperature and light conditions (from the studies by Durufle et al., 2017). Z-scores of TPM values were calculated across the samples. Yellow indicates relative up-regulation and blue indicates down-regulation.

**FIGURE S8.**
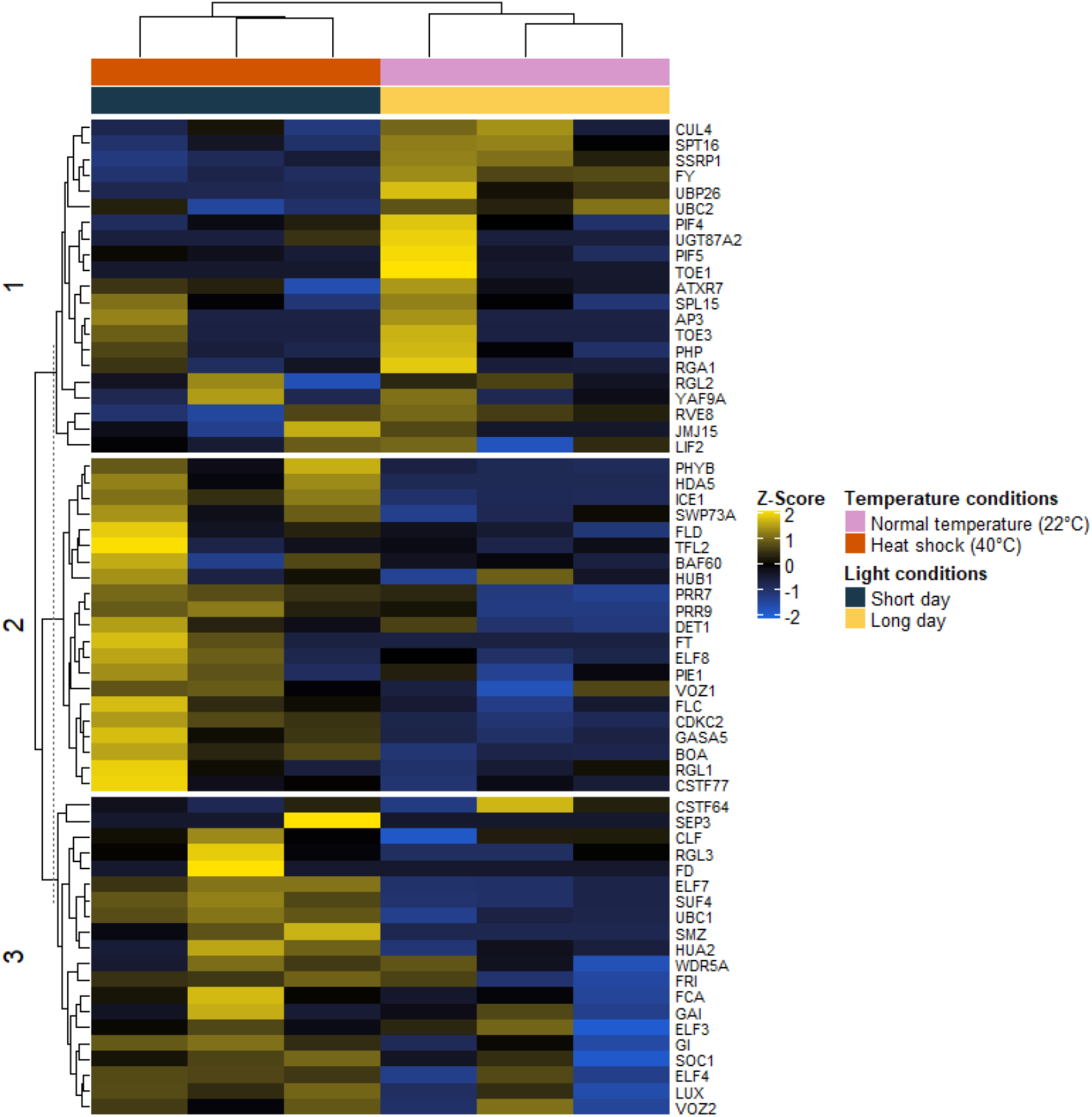
Transcriptional patterns of flowering genes in mature pollen grain. Heatmap showing the transcriptional patterns of 62 flowering-time-relating genes, namely *PIF4*, *PIF5*, *ELF3*, *FLC* and their immediate up- and downstream genes from mature pollen grain of *Arabidopsis thaliana* ecotype Col-0 grown under different temperature and light conditions (from the studies by Rahmati Ishka et al., 2018). Z-scores of TPM values were calculated across the samples. Yellow indicates relative up-regulation and blue indicates down-regulation.

**FIGURE S9.**
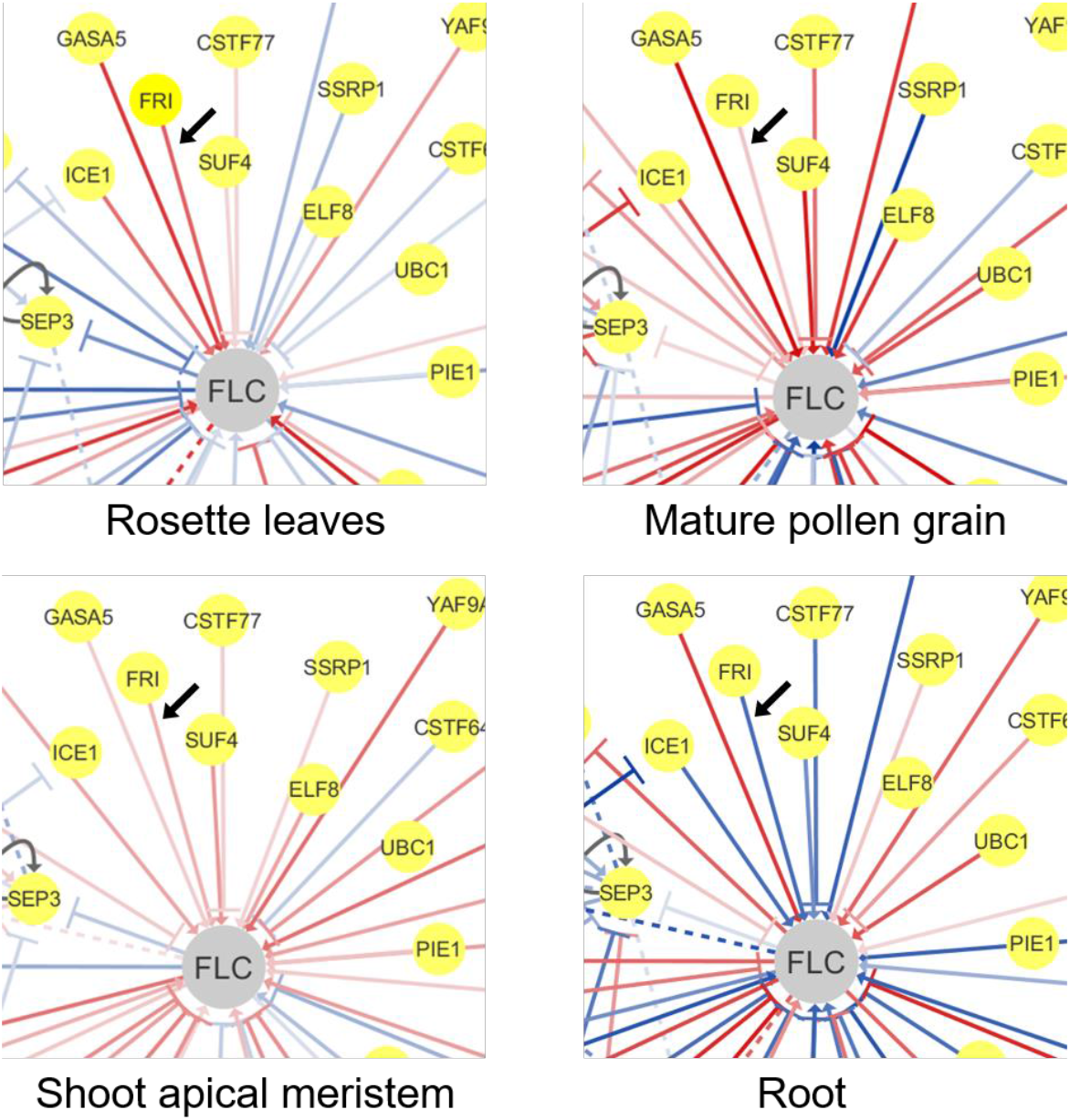
The FRI might positively regulate the *FLC* expression in tissue-specific manner. The networks showing the expression correlations which were calculated from individual tissues namely, rosette leaves, mature pollen grain, shoot apical meristem (SAM) and root. Edge colors represent the correlation coefficients: positive (red) and negative correlation (blue), using Spearman’s correlation coefficient method. Lower correlation values are shown as paler-color edges and higher correlation values as darker-color edges. Solid/dashed black lines represent the auto-regulation. Black arrows indicate the gene pair of interest.

